# Training Generalized Segmentation Networks with Real and Synthetic Cryo-ET data

**DOI:** 10.1101/2025.01.31.635598

**Authors:** Carson Purnell, Jessica Heebner, Linh Nguyen, Michael T. Swulius, Ryan Hylton, Seth Kabonick, Michael Grillo, Stephanie Grillo, Sergei Grigoryev, Frederick A. Heberle, M. Neal Waxham, Matthew T. Swulius

## Abstract

Deep learning excels at segmenting objects within noisy cryo-electron tomograms, but the approach is typically bottlenecked by access to ground truth training data. To address this issue we have developed CryoTomoSim (CTS), an open-source software package that builds coarse-grained models of macromolecular complexes embedded in vitreous ice and then simulates transmitted electron tilt series for tomographic reconstruction. Using CTS outputs, we demonstrate the effects of key microscope parameters (dose, defocus, and pixel size) on deep learning-based segmentation, and show that including both molecular crowding and diversity within synthetic datasets is key to training cellular segmentation networks from purely synthetic inputs. While very effective as initial models, the accuracy of these networks is currently limited, and real cellular data is necessary to train the most accurate and generalizable U-Nets. Using a co-training approach, we first segment over 100 tomograms from neuronal growth cones to quantify their cytoskeletal distributions and then we build a generalized cellular cryo-ET segmentation network called NeuralSeg that can segment a subset of cellular features in tomograms from all domains of life.

## Introduction

Cryo-electron tomography (cryo-ET) allows the direct three-dimensional imaging of purified macromolecules, enriched organelles, whole bacterial and archaeal cells, and eukaryotic cellular compartments in a frozen-hydrated state ^1–10^. Cryotomograms are rich with information across length scales spanning a few angstroms (Å) to many microns ^11–13^ and extracting all of this information automatically (if at all) is challenging given the variation in their content and complexity. Traditionally, human experts have been burdened with hand segmenting tomograms for both 3-dimensional (3D) representation and analysis, and this process usually takes hours to days depending on the content contained within the tomogram.

In recent years, deep learning has emerged as a powerful tool for cryo-ET image segmentation and particle picking ^14–17^, primarily due to the advent of convolutional neural networks for pixel-based pattern recognition. The U-Net ^18^ is currently one of the standard architectures for biomedical image processing ^19^ for its ability to encode feature sets within noisy inputs at a range of scales and resolution. While U-Nets are good at segmenting cryo-ET data, typically, large hand-annotated datasets are required for adequate learning and validation for generalization.

There is a rich history of simulating images in cryo-EM ^20–22^, so generating pre-annotated synthetic training data is an appealing solution for deep learning. It would allow users to quickly generate new mixtures of molecules from existing structural models and explore a range of imaging parameters, all in a matter of minutes. To this end, three cryo-ET simulators have recently been released, specifically aimed at generating training inputs for neural networks ^23–25^. FakET ^23^ uses neural style transfer to quickly generate synthetic tilt series on the scale of full modern electron detectors, but it does not provide a foundation for modeling complex mixtures of macromolecules for use as inputs by the user. Using the SHREC dataset ^26^, however, the authors demonstrated that outputs from their workflow could be used to train the U-Net-based particle picker DeepFinder ^16^. Polnet ^25^, on the other hand, coarsely models aspects of cellular ultrastructure (membrane curvatures and curved filaments) as well as mixtures of individual molecules from filtered PDBs, but does not simulate the physics of image formation in the TEM. Despite this, the authors were able to train a U-Net capable of segmenting the major constituents of real cellular tomograms, highlighting the importance of recapitulating the crowded cellular environment during modeling.

We developed CryoTomoSim (aka CTS), a full pipeline for building coarse-grained molecular models derived from any mixture of atomic coordinate files (.CIF and .PDB), alongside reconstructed tomograms based on user-defined microscope parameters and a perfect ground truth annotation. CTS provides the ability to divide individual macromolecules into submolecular annotations and supports further functionality for modeling complex macromolecular structures in various states of assembly. Additionally, CTS can generate spheroidal vesicles with embedded transmembrane and membrane-associated proteins. Users have easy control of these modeling and simulation parameters through either graphical or command-line interfaces that allow one to quickly generate a wide variety of training data.

In this paper, we document CTS (see Supplemental Text for details) and use its outputs to characterize the impact of tunable microscope parameters on U-Net segmentation accuracy, showing that training a single network on inputs from diverse simulation parameters yields a model that performs equally well on all inputs. We then go on to demonstrate the critical importance of simulating molecular crowding and diversity while training networks to segment accurately within densely packed cellular tomograms. Finally, we demonstrate that by leveraging the convenience of synthetic data to train “seed networks”, one can quickly train a generalized U-Net through the iterative addition of relatively small amounts of hand-corrected real data.

## Results

### Training U-Nets with CTS Outputs

First, we developed a streamlined workflow for training segmentation U-Nets directly from CTS outputs (Figures 1 and 2). The workflow requires no hand annotation and can be easily set up in minutes within the Dragonfly deep learning environment, or any environment of your choosing. As a first challenge, we sought to segment *in vitro*-assembled “cofilactin” filaments that were plunge-frozen in a thin sheet of polymerization buffer. Because the tomograms included regions of cofilactin and bare F-actin, we used both filament types in our synthetic dataset. To construct the training set, we helically extended short segments of cofilactin (PDB ID 3J0S) and F-actin (PDB ID 6T1Y) to three times their original length (Figure 1A) within Chimera. Cofilactin was split into separate submodels containing the actin filament (red) and cofilin monomers (blue), and these submodels were grouped and saved as a single PDB appended with “.complex.pdb”, so that CTS would label the submodels as separate classes in the ground truth atlas.

**Figure 1:**
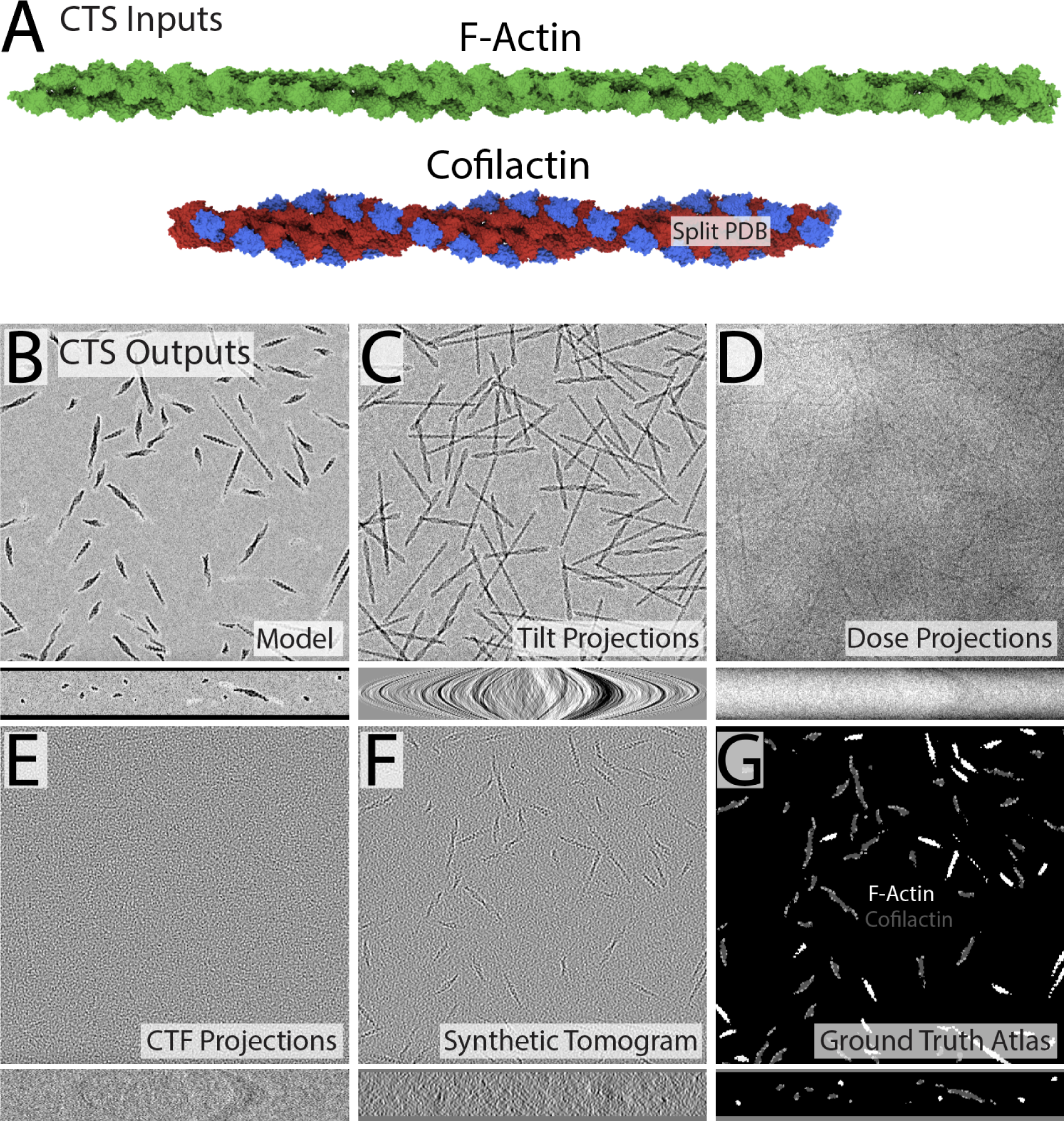
CTS workflow. (A) Space-filling representation of F-Actin (green, PDB 6T1Y) and cofilactin (PDB 3J0S) split into its constituent F-actin (red) and cofilin (blue) components, which were used as input to CTS for model building. (B-G) Representations of all the outputs from CTS, which were generated in modeling and simulation steps that took less than 5 minutes combined. (B) Volumetric 3D model of F-actin and cofilactin filaments placed randomly within a 400x400x50 voxel volume at 12 angstroms/voxel. The upper and lower bounds were constrained so that only filaments lying nearly flat were accepted, simulating the geometry of the air-water interface during plunge freezing. The top panel shows a slice through the XY axis and the lower panel shows a slice through the XZ axis. (C) The model from (B) was used by CTS to calculate a dose-symmetric set of “Tilt Projections” from +60 to -60 every 2 degrees. Upper panel: Zero degree tilt projection. Lower panel: View down the XZ axis of the full tilt series. (D) Tilt series with electron dose applied. (E) Tilt series with 2.5D CTF convolution further applied. (F) The dose/CTF-corrupted tilt series was used by IMOD to automatically calculate a synthetic tomogram. (G) A “ground truth atlas” was also generated by CTS, which contained a perfect per-voxel annotation of the two PDB inputs, including the cofilin submodels.

**Figure 2:**
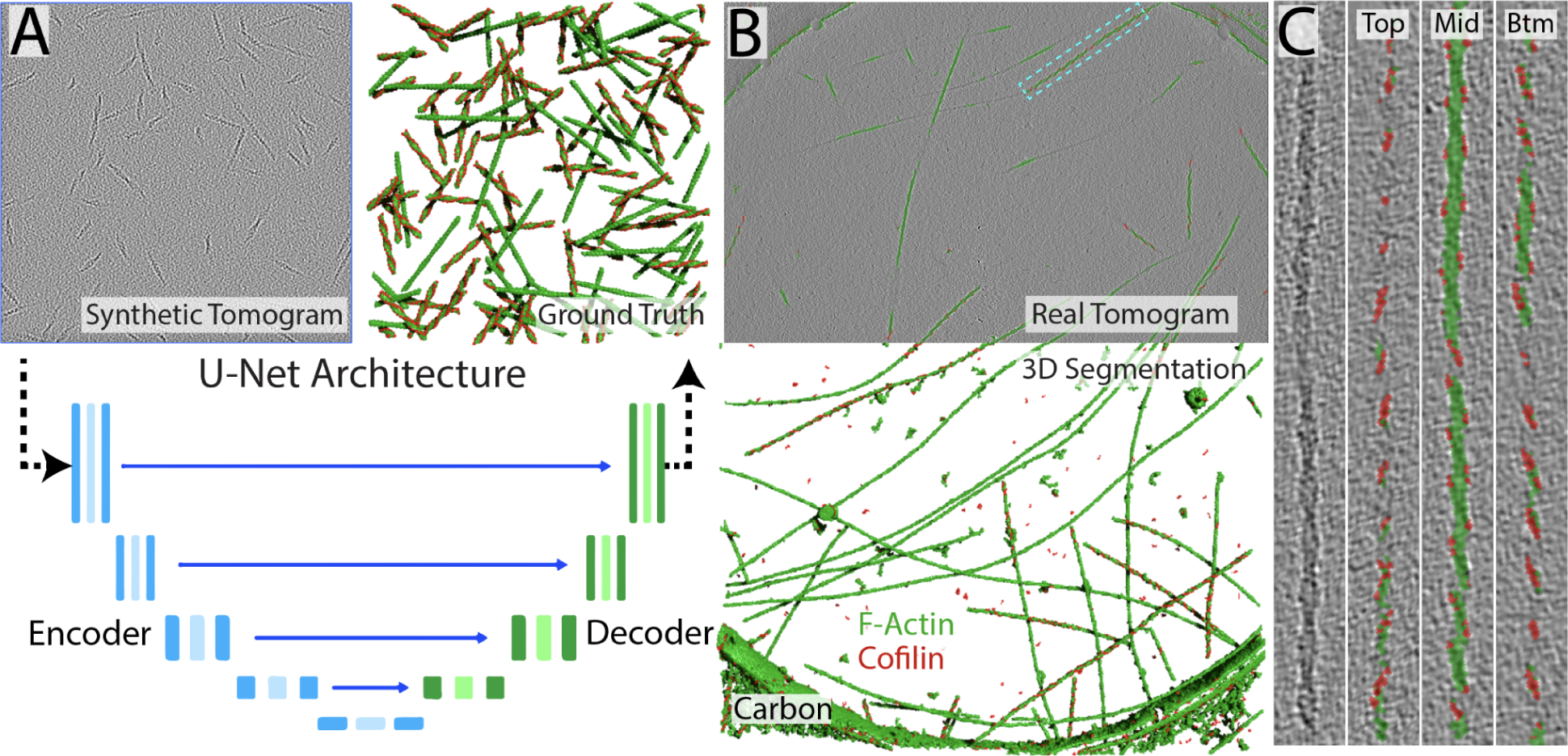
Deep Learning-based segmentation workflow. (A) Schematic of the information flow through the U-Net during training. The top left panel is the synthetic tomogram generated by CTS, which is fed as 2D patches through the encoder/decoder arms that form the classic U-Net architecture, and the top right panel is the CTS-generated ground truth atlas used for training. (B) Inference of the trained network to real tomographic data containing a mixture of F-Actin and cofilactin filaments. The top panel shows the segmentation overlaid on the tomographic data directly and the bottom panel reveals the full 3D segmentation. The edge of the carbon hole can be seen in both panels. (C) Zoom-in and sectioning through a single cofilactin filament that is boxed in panel (B). Slices are through the top, middle and bottom of the same filament, revealing the right-handed helical turn of the cofilins bound to the actin filament.

Next, we gave these two PDBs to CTS as inputs and asked it to generate a 400x400x50 voxel model at 12 Å/pixel with 400 iterations of filament placement. To approximate the air-water interface during sample blotting, constraints were added to the upper and lower bounds of the Z-axis so that only filaments falling completely within these bounds were considered for placement. If the filament was free from overlap with any other filaments, it was accepted. Many placements were rejected, but the accepted subset of filaments were all lying relatively flat with respect to the XY plane (Figure 1B), similar to real tomograms of plunge-frozen cofilactin filaments. As a final step, a field of vitreous “water” was added to the background.

The model generated in the previous steps was then used by CTS to calculate a dose-symmetric set of tilt projections every 2 degrees from +60 to -60 degrees (Figure 1C). This tilt series was corrupted by electron dose (100 total e^-^/Å^2) and a contrast transfer function (defocus: -4 µM) before automatic reconstruction via weighted back projection into a synthetic cryotomogram via IMOD ^12,27^ (Figure 1D-F). Finally, CTS generated a fully annotated ground truth atlas of the voxels belonging to F-actin and cofilactin (Figure 1G).

The synthetic tomogram and its accompanying ground truth were imported into Dragonfly, where we merged the F-actin classes from both filament types into one F-actin class (green) and one cofilin class (red), and then used this remapped ground truth to train a multiclass U-Net (Figure 2A). By 70 epochs the network had converged (Movie S1), and was then inferred to a real cryotomogram containing *in vitro* actin and cofilactin filaments (Figures 2B & C). The segmentations are imperfect in that not every individual cofilin is delineated, but the right-handed helical path of the cofilin signal is striking. It is a level of detail that almost no human expert could achieve in the minutes it took for the U-Net to process the whole tomogram.

To further test our workflow on various macromolecular shapes and textures, we applied the same approach to Calmodulin-dependent kinase II (CaMKII) holoenzymes, proteasomes, and condensed human chromatin particles. In each case, networks trained only on CTS-generated datasets were able to segment different protein regions or subunits with high accuracy (Figure S1). With CaMKII, there were small clusters of holoenzymes that were ignored because such clusters were not represented in the training set. Additionally, not every visible kinase domain is segmented because in reality they are highly flexible compared to the static model used for training. In the chromatin arrays, the DNA and histone components of the individual nucleosomes are well segmented, but the DNA linkers stringing them together are only partially captured. This is due to the fact that the ordered linkers in the PDB model used to train the network do not match the variable linker lengths and conformations observed in human heterochromatin ^28^. That being said, these results constitute an exciting leap forward in terms of the ease with which the granular segmentations necessary for quantitative analysis can be achieved. Moving forward, our plan is to develop methods to better mimic real molecular flexibility in our initial CTS models to increase the efficiency of *in vitro* segmentation networks.

### Exploring Microscope Parameters

The primary advantages of building a cryo-ET simulator are that you can easily alter imaging parameters without spending money or time at the microscope and, on top of that, you have a calculated ground truth to compare your segmentations to. Because of this, we explored the robustness of U-Nets to the microscope parameters typically varied during data collection: electron dose, defocus, and pixel size. We started with total electron dose (e^-^/Å^2^ per tilt series) because it has a direct impact on the available signal in cryotomograms. To do this, we locked the defocus to -5 microns and the pixel size to 10 Å/pixel, before creating a series of synthetic tomograms (between 1 and 100 e^-^/Å^2^) from a model of randomly oriented ribosomes (Figure S2A). Ribosomes were chosen for their abundance in cellular tomograms, as well as their asymmetric nature. The goal of the experiment was to test the robustness of U-Nets trained with a single-dose input when inferred to tomograms simulated at other electron doses.

As one might expect, the network trained at the highest dose (100 e^-^/Å^2^) was the most accurate in terms of DICE scores when there was sufficient signal (30 e^-^/Å^2^ or above), otherwise, the perceptural fidelity fell off sharply as the dose dropped below 10 e^-^/Å^2^ (Figure S2 B&E). The network trained at the lowest dose (1 e^-^/Å^2^) robustly detected and segmented all ribosomes at all signal levels teste (Figure S2 C), but never as precisely as the high-signal network under ideal conditions according to DICE scores (Figure S2 E&F). Finally, we found that by training a network with both low- and high-signal inputs simultaneously, a model emerged that captured the accuracy and sensitivity of both individually trained networks (Figure S2 D&E). While the effect of defocus on deep segmentation was more modest than electron dose, the same general theme held true, with a falloff in DICE scores as you deviate from the defocus value used to train the U-Net (Figure S3), and training at multiple defocus values helped to mitigate defocus-mismatch effects during segmentation.

Pixel size has a similar impact on segmentation efficiency in that moving too far from your originally trained pixel size leads to suboptimal segmentations. In our case, a network trained at 10 Å/pixel was inferred to simulated tomograms of ribosomes across a range of pixel sizes from 5 Å/pixel to 15 Å/pixel. What we saw is that when the pixel size reaches half that of the training dataset (5 Å/pixel), individual instances of ribosomal segmentation begin to break apart, visually (Figure S4 A&B), and the DICE score also begins to drop (Figure S4 D). As pixel size moves in the other direction (15 Å/pixel), the DICE begins to fall again as individual ribosomes become too small to be recognized by the network (Figure S4 C&D).

While these results mostly seem intuitive, these experiments helped to establish guidelines for how these common microscope parameters are tolerated by U-Nets. They suggest that the signal available in the tomogram and pixel size are the primary factors in determining performance of a network during segmentation. That said, networks trained at a single pixel size were still surprisingly robust to deviations as long as they did not stray too far.

### Segmenting *In Situ* Tomograms

Next, we chose a neuronal tomogram densely packed with microtubules and tested our approach to segment them within crowded cytoplasm. A U-Net trained only on synthetic microtubule fragments failed to cleanly segment the microtubules within the real tomogram (Figure 3A). Given the crowded nature of cytoplasm, we reasoned that adding other protein components to the training data as “distractors’’ would lead to improved performance by the network, which turned out to be true (Figure 3B&C). What we learned was that increasing the packing density of ∼15 small crowding agents (<200 kDa) had a significant effect, but with diminishing returns as packing density was increased. By replacing the distractor set with ∼15 larger distractors (>200 kDa), we were able to see further increases in performance, suggesting that different layers of crowding agents might be important for network performance. Unsurprisingly then, our best network was trained on a mixture of larger macromolecules that were layered over with a dense packing of smaller macromolecules to a high density within the synthetic tomogram. The gradient of improvement from increasing crowding and molecular complexity is clearly observed in Figure 3B.

**Figure 3:**
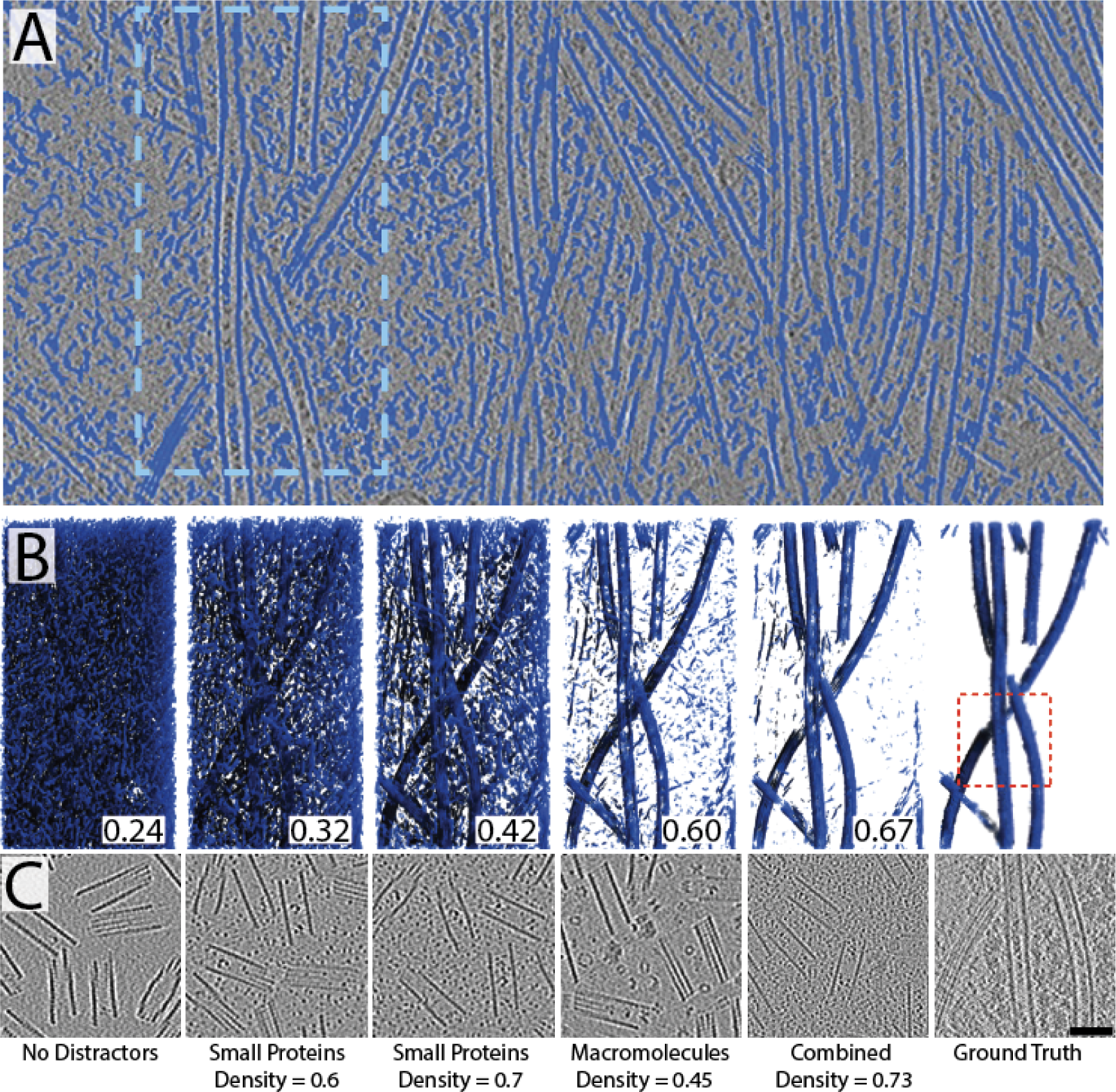
Segmenting cellular data with synthetically trained U-Nets requires increased crowding and complexity. (A) Slice through a poorly segmented neuronal tomogram dense with microtubules running vertically. It is clear that the microtubules were segmented along with nearly every other protein density (blue). The light blue dashed-box represents the region depicted in (B) for comparison of network performance. (B) Panels (from left to right) depicting 3D segmentations of the region boxed out in (A) by networks trained on the synthetic data shown in (C). The numbers in (B) are DICE scores calculated based on their similarity to the ground truth annotation shown in the far right panel.

Given the success of this strategy, we trained a four-class U-Net to segment membranes, microtubules, actin filaments, and ribosomes simultaneously within crowded “synthetic cytoplasm”, which included multiple heavy layers of distractors. The results of inferring the same network to four disparate *in situ* tomograms are shown in Figure 4. While the segmentations have errors (some MTs were misclassified as actin in Figure 4C and mitochondrial membrane is misclassified as actin in Figure 4D), they are quite remarkable given that no expert annotation was necessary to set up the training inputs. Panels 4A & C show segmentations from two different neuronal tomograms, while panels 4B & D show segmentations of FIB-milled lamella from both a HeLa cell (EMDB-11992) and a *C. elegans* embryo (EMDB-4869), respectively. For a better sense of network performance we suggest you see movie S2, where the segmentations are overlaid onto the tomograms in all three dimensions. While networks trained solely on synthetic data show much promise, under these conditions, they do not yet capture the full variability observed in real cellular tomograms. They do, however, function as a great starting point for developing better models with the help of real data, as we’ll see below, and they can greatly accelerate the segmentation of a rough ground truth for hand-correction and continued training.

**Figure 4:**
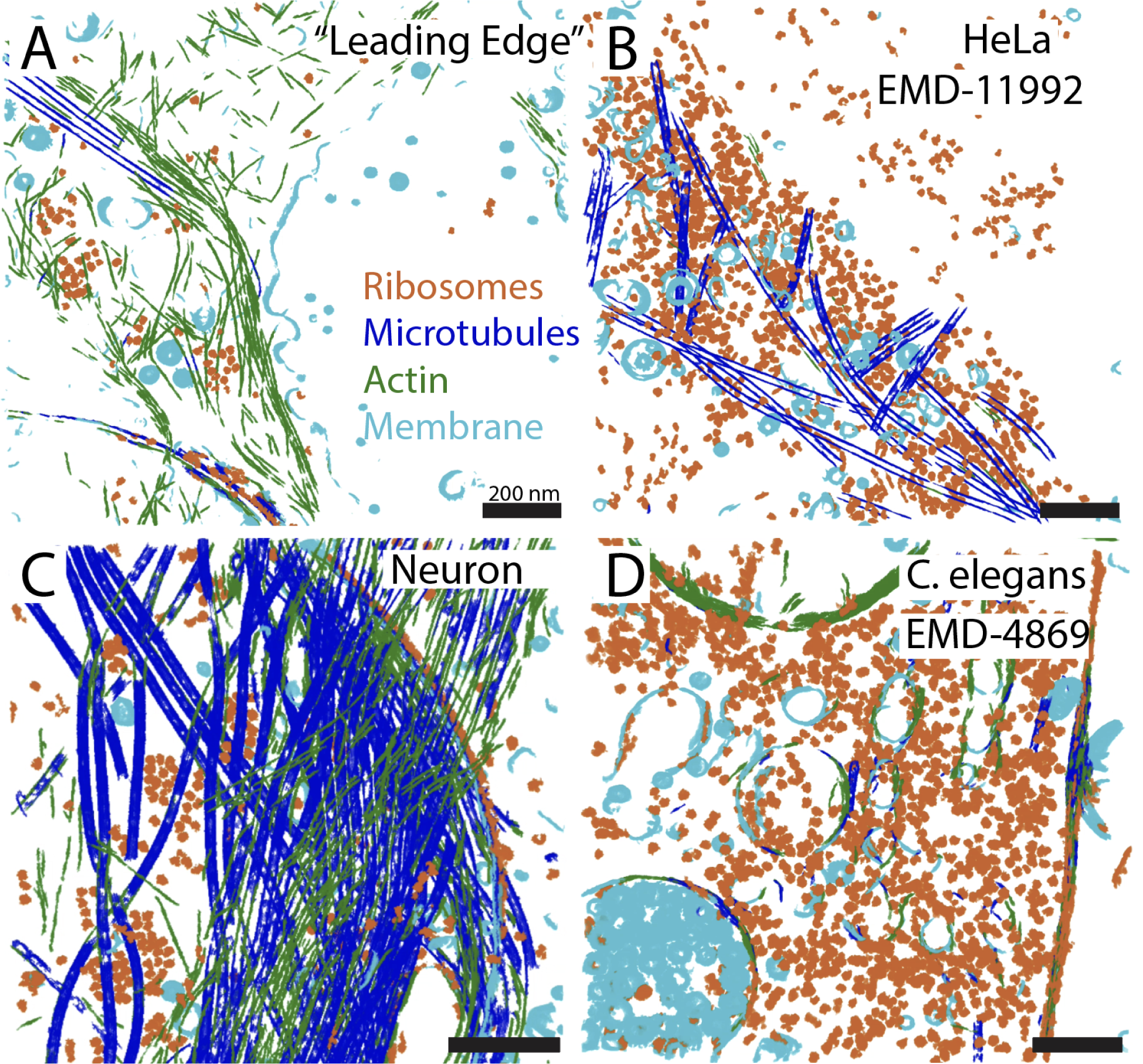
Networks trained on synthetic cytoplasm alone are clean but objects are mis-classified. A single U-net trained solely on synthetic cytoplasm was used to segment (A) a neuronal tomogram at the leading edge of the cell, (B) a FIB-milled HeLa cell, (C) another neuronal tomogram rich with microtubules, and (D) a FIB-milled *C. elegans* embryo. See movie S2.

### Co-Training with Real and Synthetic Data

We reasoned that adding real cellular data and context to our training sets would increase their accuracy and generalizability across tomograms, but we wanted to use minimal amounts of hand-annotation by human experts. To do this, a 400x400x50 voxel subvolume from the “leading edge” segmentation in Figure 4A was selected for its balance of microtubules, actin filaments, membrane, and ribosomal content (for this experiment, F-actin and cofilactin were merged into a single actin class for simplicity). The segmentation was expertly corrected through class remapping and removal of false positives/negatives within Dragonfly. This small hand-corrected segmentation was then considered to be ground truth (Figure 5D) in an experiment designed to look for synergy between real and synthetic inputs.

**Figure 5:**
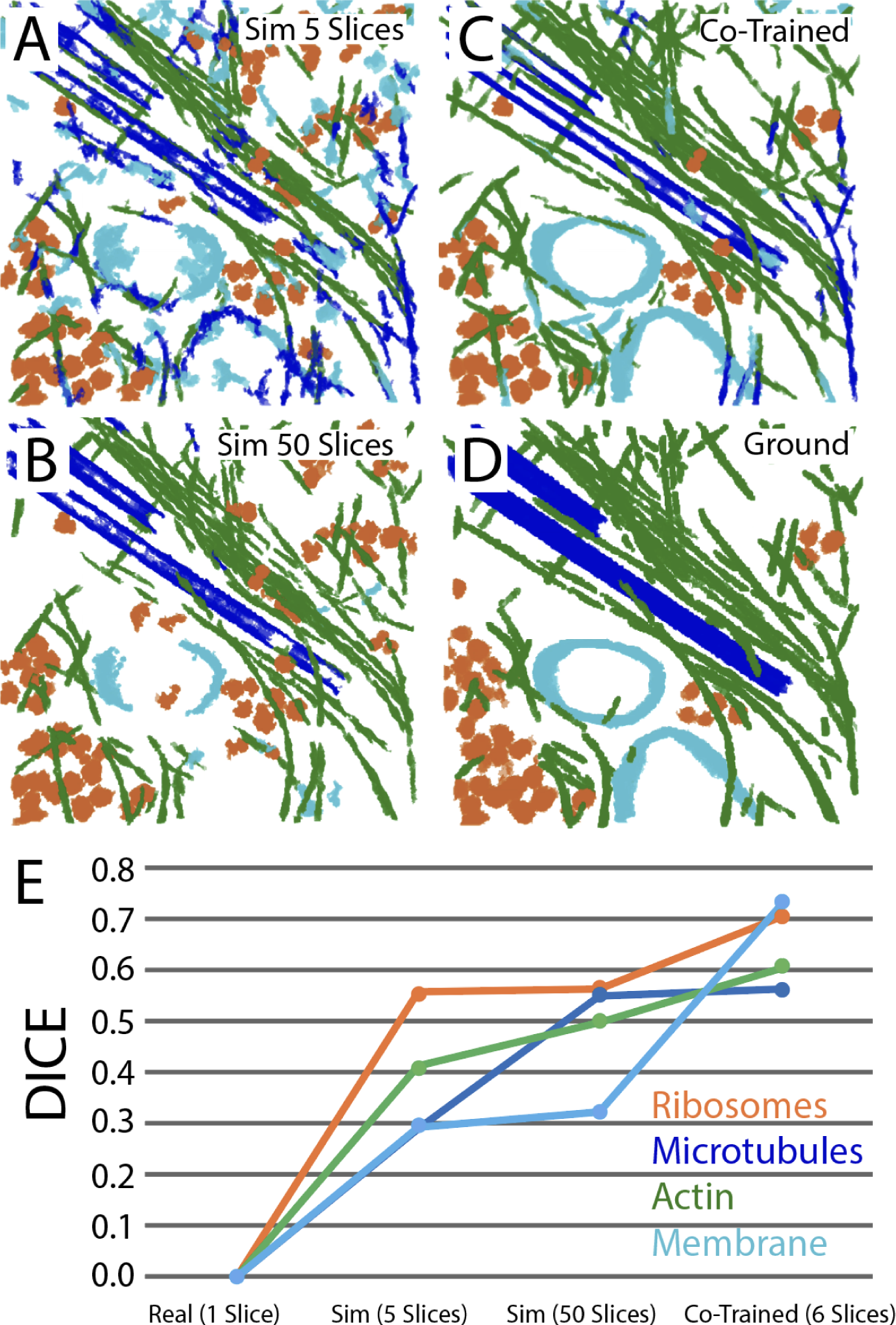
Augmenting simulations with hand annotations of real data. 3D segmentations of a 400x400x50 voxel slab of the “leading edge” tomogram from Fig. 4A using networks trained on either (A) five slices of simulated cytoplasm, (B) 50 slices of simulated cytoplasm, or (C) co-trained on the same five slices of simulated cytoplasm plus 1 slice of the ground truth in (D). (E) DICE scores for each class, calculated for each network.

To start, we trained a U-Net on a single central slice from the ground truth (Patch size = 128x128x1, stride ratio = 1, batch size = 8). Under these conditions, the input slice of 400x400x1 failed to train a functioning network, and the DICE score for each class was zero (Figure 5E). Next, networks trained on either 5 or 50 slices of synthetic cytoplasm were compared (Figure 5A,B&E). Both perceptually and according to DICE scores, 50 slices outperformed 5 slices, though the improvements did not scale proportionally to the input data. Interestingly, by adding the same single slice of ground truth data to the 5 slices of synthetic data and co-training the U-Net, a network emerged that outperformed the 50 slices of synthetic data on all classes (Figure 5C&E).

Next, to test the utility of co-training to deal with large datasets and to push the limits of our approach, we added the additional challenges of discriminating between actin and cofilactin *in situ* (Figure S5), as well as the protein folding chaperone TriC, a multimeric complex often found intracellularly alongside ribosomes ^12^. For this task, 106 cryotomograms of E18 rat hippocampal growth cones were compiled from four years of previous data collection. Each was assigned to a growth cone region by referencing the lower magnification search image and identifying its relative location within the morphology of the growth cone (Figure 6A). Peripheral (P)-domain was assigned whenever a tomogram was clearly at the edge of a cell over a filopodia or the lamellipodial veil. Central (C)-domain was assigned to tomograms that were positioned in the middle of the growth cone, and Transition (T)-zone was assigned to tomograms that fell between the P-and C-domain. In total, the dataset spans twenty individual growth cones where 23 tomograms were assigned to the C-domain, 27 to the T-zone and 56 to the P-domain.

**Figure 6:**
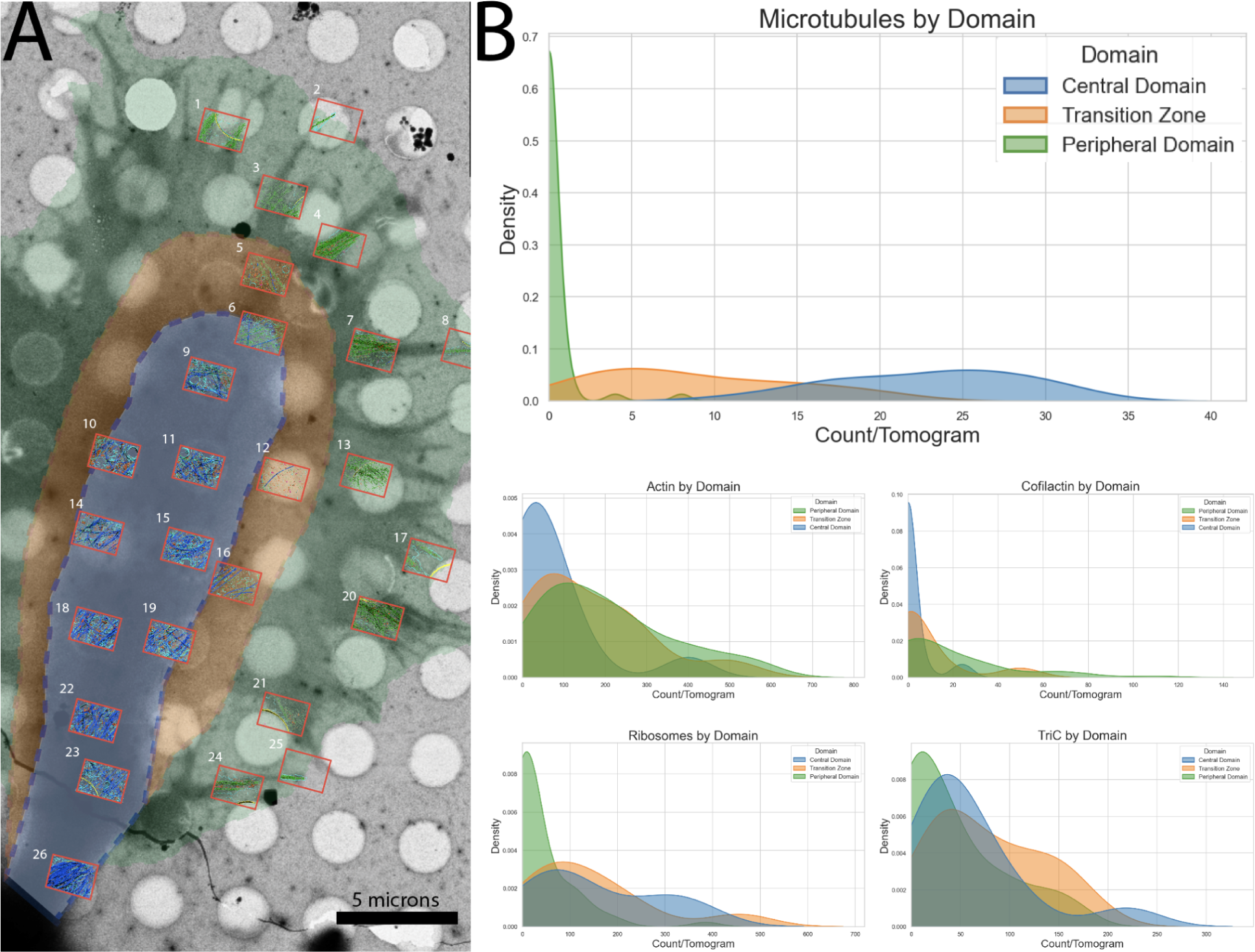
Quantitative segmentation of 106 neuronal growth cone tomograms. (A) Domain-colorized low-magnification overview of a vitrified growth cone on a holey carbon grid, overlayed with scaled segmentations from 26 individual tomograms. (B) Smoothed histograms of particle class abundances, within individual tomograms, from different growth cone domains (Central:blue, Transition Zone:orange, and Peripheral:green). For Microtubules, ribosomes and TriC, count represents actual particle count, but for actin and cofilactin, the total voxel count within each tomogram was used.

We then trained a U-Net to segment nine classes simultaneously (membrane, microtubules, actin, cofilactin, ribosomes, TriC, the carbon edge, gold fiducials, and background). This network was co-trained on two datasets then applied to all 106 tomograms. The real training data was a 600x600x55 hand-segmented cropped section of the “leading edge” tomogram containing all nine classes. The synthetic dataset was a 400x400x50 simulated tomogram containing the same nine classes embedded in dense distractors. At several instances during segmentation, the network was copied and the duplicate was further trained with additional real tomographic data from the trouble datasets. Typically this was because the dataset encountered was of poorer quality in some way. Each retrained network, however, allowed the set to segment features across a broader range of imaging conditions. It took training a total of three networks to segment all 106 tomograms, and when that was finished, islands of noise smaller than 50 connected voxels were removed before quantification (see methods for counting procedures).

Within the domains of the growth cone, subpopulations of known cytoskeletal composition have been shown to exist by cryo-ET ^29^, making them ideal for assessing the performance of our networks on a cellular scale. It is known, for instance, that the P-domain is composed almost entirely of actin, while the T-zone is a mixture of pioneering microtubules and actin, and the C-domain is enriched in microtubules and membrane-bound organelles ^29^. Growth cone domains are colorized on the overlay in Figure 6A, and when cytoskeletal/ribosomal content was measured across these domains using our segmentations, we observed very similar distributions to those reported by Atherton et al (see Figure 6B).

Microtubules (932 total) were nearly absent from the peripheral domain and densely packed in the central domain. F-actin was present in nearly all (97) tomograms and of those, 56 also contained some amount of cofilactin. While less pronounced, the gradient of actin/cofilactin runs in the opposite direction of microtubules (denser at the periphery), as expected from previous publications from our lab ^30^. It is worth noting that all counts in Figure 6B are presented as fully instantiated molecules/tomogram except for the “counts” of actin and cofilactin, which are presented as total voxel count/10,000. This is because the thin, densely packed filaments are not trivial to instantiate and are the subject of further methods under development in the lab. While having an accurate filament count, along with their length measurements, will bring new insight to the structure of actin networks in the growth cone, it should not fundamentally change the spatial distribution of actin and cofilactin across these domains. 12,759 ribosomes were segmented in total, including some small clusters of closely grouped ribosomes that were not easily separated algorithmically, and 7717 TriC particles were identified based on watershed analysis of the segmentations. Given the accurate instantiation of individual TriC molecules, clustering analysis was performed and it revealed TriC’s tendency to form small clusters (∼5 molecules) throughout the cytoplasm. In all 84 tomograms where TriC was identified, there was at least one cluster (Figure S6A). While TriC was clustered more frequently in all regions, no statistically significant effect of region, on clustering, was detected (Figure S6B). The mean number of TriC proteins per cluster was relatively consistent across all domains (Fig S6C), and the mean nearest neighbor distance between particles in a cluster was as well (Figure 12D), confirming the tight intra-cluster packing observed in the tomograms (Figure S6A). Independent of their clustering, ribosomes and TriC are more evenly distributed across all domains of the growth cone than the cytoskeleton. Due to the region-dependent variability of cytoskeletal proteins in the growth cone, this data suggests ribosomes and TriC are likely more free to move about the crowded cytoplasm.

### Iteratively Co-Training a Generalized Network

Real cryo-ET datasets do not come with ground truths to facilitate co-training, but “seed networks” trained primarily on synthetic data can quickly segment real data to be hand-corrected and used as a ground truth. We decided to leverage a seed network strategy to begin iteratively co-training a single, more generalized cellular segmentation U-Net. To do this we first established a set of 10 tomograms, derived from synthetic cytoplasmic models, to be used in all iterations of network training (Figure S7A). These were all 400x400x50 voxels, generated programmatically by CTS to vary randomly within a set range of modeling (pixel size and molecular crowding) and simulation parameters (dose and defocus). All individual synthetic tomograms contained the primary targets (membrane, microtubules, actin, cofilactin, ribosomes and TriC) as well as randomized sets of 10 distractor molecules (5 large and 5 small) from a pool of 20 possible distractor PDBs for added cytoplasmic variation.

Next, we co-trained a new base network on our block of normalized and concatenated synthetic data, along with a small amount of real data from the leading edge tomogram (Figure S7B, 1.1). We inferred this base network to approximately 10 varied neuronal tomograms and examined them for regions where the network struggled. After identifying several areas, small subvolumes (400x400x∼20 voxels) of the tomograms and their accompanying segmentations were cropped and corrected before normalization and concatenation for re-training of the network (1.2). This time, for each subiteration of training, everything in the previous training sessions was included and a brand new network was trained.

In iteration 2.0, we repeated the above process using a set of three non-neuronal tomograms (Figure S7B, 2.1). We segmented them with our best network, hand-corrected subvolumes from them, and concatenated them into our real training data. For iteration 3.0, we focused on eliminating common tomographic features that are often mis-classified (ice-surface contamination, fiducial gold, carbon surface and edges, and tomographic fringing). To do this, we cropped subvolumes containing these features, merged all the mis-classified pixels into the background class, and concatenated them to our block of real data (Figure 12B).

The final result is NeuralSeg, a generalized U-Net for the segmentation of basic cellular structures in cryo-ET data (Figure 7). To explicitly assess the limitations of the network, we ran NeuralSeg through a battery of experiments. First, pixel size was tested by rescaling a neuronal tomogram that was generated at 10 Å/pixel to 20 Å/pixel in 20% increments, and running each of them through NeuralSeg (Figure S8). At first glance, the results look surprisingly similar across the entire range, but there was a falloff in performance at the two extremes. At 10 Å/pixel, there was granular noise visible in the segmentation, which was generally mis-classified as ribosomes (orange). The TriC class (purple) was sparsely populated at this pixel size and the microtubules (dark blue) were only segmented near their central slices, as well as often misclassified as membrane (light blue). At 20 Å/pixel, the granular noise was nearly gone but so were many of the particle instances identified at the middle range, along with quite a few individual actin filaments (green). By visual inspection, the segmentations at 12 and 14 Å/pixel are the most complete, and this is interesting given this is precisely the size-range represented in our simulated block of training data (Figure S7A).

**Figure 7:**
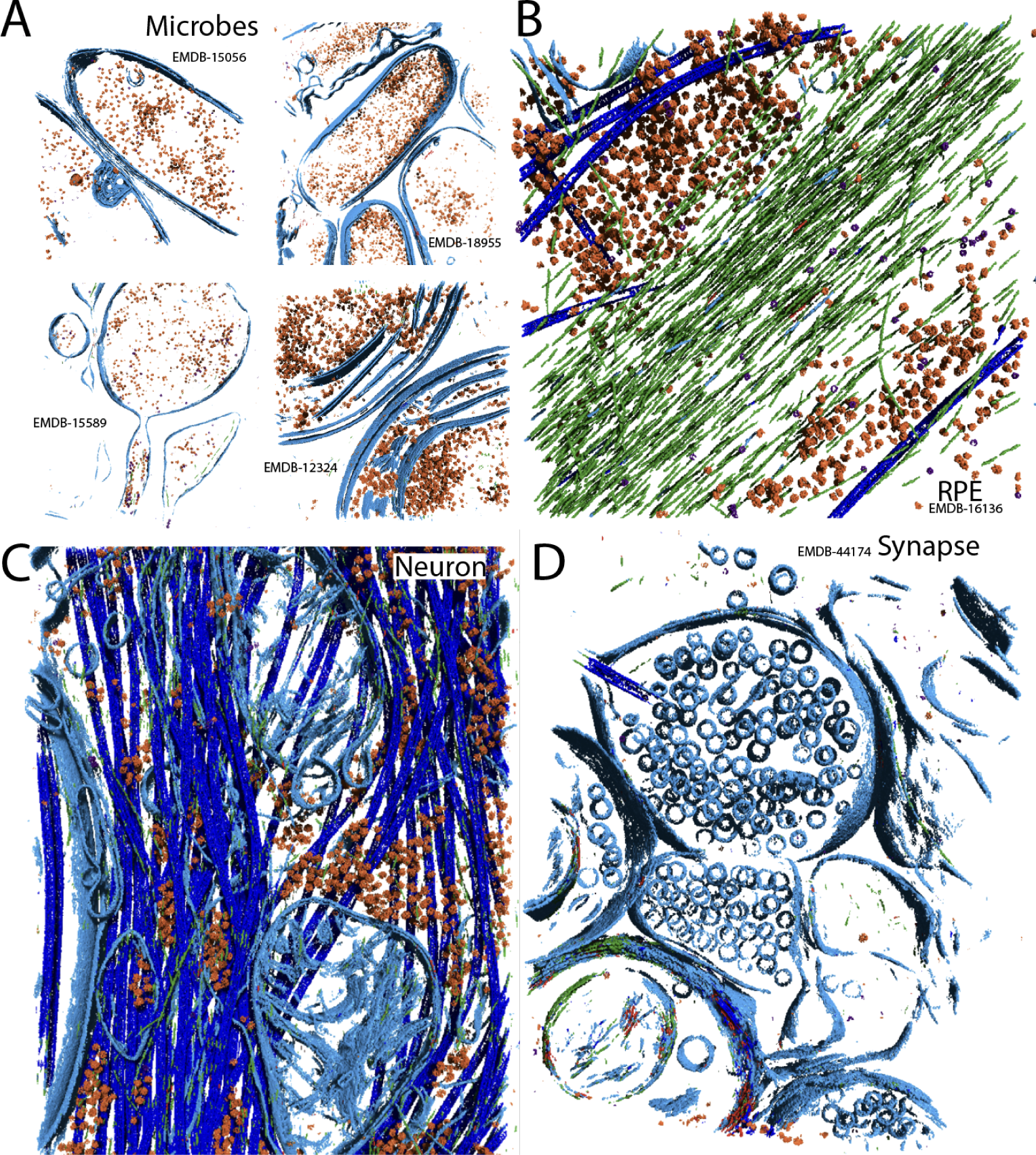
NeuralSeg can segment most cellular tomograms. The NeuralSeg U-Net was used to segment (A) four microbes of varying shape, (B) a lamella milled from an RPE cell, (C) an intact neuron adhered to the grid, and (D) a glutamatergic synapse milled from vitrified hippocampal tissue.

Next, we tested the network’s ability to segment tomograms from -4 to -10 µM defocus (Figure S9) using tomograms collected along neurites in the same grid square region displaying uniform contrast, to minimize differences from sample thickness. Given U-Net robustness to defocus (Figure S3), we were not surprised to see the network performed across the entire range. It is worth noting that much of the real data used in training NeuralSeg was collected at defoci lower than -6 µM. In fact, there is no real data in the training set collected at -4 µM defocus, which might explain why the network appeared to struggle a bit at this defocus according to the “fullness” of the microtubule segmentations (dark blue).

Finally, we tested the network’s performance on lower contrast data, due either to tomogram thickness or low electron dose used during collection. To do this, regions from the same grid square with different sample thickness (varied mean intensity) were identified for tilt series collection with a total dose of 180 e^-^/Å^2^ (the typical dose used in previous experiments). Unsurprisingly, the thickness of the samples was inversely correlated with the performance of the network. For instance, at ∼250-nm thick, NeuralSeg still performed reasonably well (Figure S10A) but under the same beam conditions, ∼300-nm thick leads to obviously reduced contrast and segmentation performance (Figure S10B). By 400-nm thick, the network almost fails to segment entirely (Figure S10C).

To explicitly test electron dose, we collected a tilt series at half the previous dose (90 e^-^/Å^2^) and then a second tilt series in the same spot at half that dose (45 e^-^/Å^2^). At 90 e^-^/Å^2^ the segmentation is still reasonably good but the further signal degredation at 45 e^-^/Å^2^ lead to a poor segmentation (Figure S10D&E). While this experiment introduced the risk of radiation damage due to exposing the same position to two different tilt series, the total exposure falls under 150 e^-^/Å^2^ suggesting that the network’s performance loss is based on the resulting contrast loss from lowered electron dose and not radiation damage. Taken together, these experiments help to define upper and lower bounds for NerualSeg’s reasonable performance that should match the data collection parameters for most labs collecting cellular tomograms.

## Discussion

Cellular cryotomograms are complex. Depending on the optical parameters used for data collection, and the state of the sample, a single tomogram can contain information on the micron scale down to a few angstroms ^11–13^, and a dataset might contain tens or thousands of tomograms ^31^. As montage tomography develops ^32,33^, the length scale grows, and this is being multiplied by the promise of serial cryo-liftout techniques ^34^. Developing tools to parse the biomolecular structures within these 3D maps is paramount to understanding cell and tissue function. To this end, we have developed CryoTomoSim (CTS), an open-source software suite that builds coarse-grained models of macromolecular mixtures (including membrane-embedded complexes). It then simulates transmitted electron tilt series for tomographic reconstruction, alongside a perfectly annotated molecular ground truth atlas. Here, we have discovered that synergy exists between real and CTS-simulated data.

Our lab and Martinez-Sanchez et al (2024) have demonstrated that the cytoplasmic landscape can largely be synthesized when training tomography-based segmentation networks, and that molecular crowding is an important part of the equation. While the results make us hopeful for a future where synthetic data could be all one needs, currently, networks trained solely on synthetic data are imperfect. Perhaps cellular modeling will advance enough for us to solve the problem by simulation alone, but until then, we believe iterative co-training will serve as an effective approach for capturing both the molecular detail of cytoplasmic components as well as the cellular context needed for accurate and generalizable predictions by neural networks.

Currently, the biggest limitation in our workflow is the need for human expertise and input. That being said, human intervention is relatively painless because it involves deciding which regions of the tomogram the network is underperforming on, and how it is underperforming, so small subtomograms can be identified and hand corrected. If we’re lucky, AI will one day make these decisions as well, but until then this approach constitutes a major advance in the user experience and workflow required for segmenting large cryotomographic datasets.

CTS was originally developed in our lab for helping train networks to better discriminate between F-actin and cofilactin, and its performance exceeded our expectations, encouraging extensions into segmenting other cytoplasmic features. We cannot predict how helpful it will be for others, but our lab has successfully used simulated data as a starting point for co-training a variety of segmentation networks. We plan to train and release networks regularly for the public, but it is our hope that CTS and the workflow illustrated here will give others the power to build better networks for their own segmentation problems.

We choose to develop models and to work within the Dragonfly environment (Comet, Inc), but it is by no means a requirement to take advantage of CTS outputs. We have trained many U-Nets directly in Python using TensorFlow ^35^, but we developed this Dragonfly-based workflow for ourselves and believe it provides the flexible, reliable, and batchable environment that many tomography labs will prefer. Additionally, deep learning is complicated and tomogram segmentation requires a robust set of visualization/annotation tools, and these are already embedded in Dragonfly. For labs developing networks outside of Dragonfly, we have made the normalized and concatenated training data available so that your own model can be trained locally and improved upon for your custom needs.

There is still a ways to go before all the important features within cellular tomograms can always be quickly segmented for analysis, but we believe that this study represents several milestones in the quest to solve the segmentation problem in cryo-ET. 1) NeuralSeg was conceived and developed from scratch in only a few days, once we found an optimal workflow. It required no hand segmentation beyond the class remapping and false positive/negative correction within relatively small subvolumes, and this correction usually did not take more than 10 minutes. Perhaps most exciting is that NeuralSeg will definitely be improved upon quickly. 2) We have successfully trained U-Nets to discriminate between bare and cofilin-decorated actin filaments within the crowded milieu of over 100 growth cone tomograms of varying thickness, and mapped the distribution of cofilin’s interaction with F-actin across subcellular domains. 3) Through iterative co-training, we have established a generalized segmentation model that can segment tomograms from all the known domains of life, prepared in all the known fashions required for cryo-ET imaging (plunge-frozen or high-pressure frozen, intact or FIB-milled). The biggest challenge, now, is to utilize the wealth of information captured by these segmentation networks in truly meaningful ways.

## Methods

### Protein Purification

Purified non-muscle actin (Cat # BK013) and cofilin (Cat # CF01-A) were purchased from Cytoskeleton, Inc. (Denver, CO). Alpha-CamKII holoenzymes were purified as described in detail here ^36^. Briefly, rat alpha-CaMKII was expressed in and purified from Sf21 cells using a Calmodulin-Sepharose column before size exclusion chromatography.

### Chromatin Isolation

Native Chromatin was isolated according to the detailed methods described in ^28^. Briefly, K562 cell nuclei were isolated and treated with micrococcal nuclease (MNase). The size of the nuclease fragments was determined by agarose DNA electrophoresis and the concentration of soluble DNA was monitored spectrophotometrically. The released soluble chromatin was concentrated to a final concentration ranging from 4.0-8.0 mg/mL.

### Neuronal Cell Culture

For neurons cultured on grids, we followed the protocol described in detail here ^37^. Briefly, primary neurons were harvested from E18 Sprague Dawley rat hippocampi (acquired from BrainBits LLC, Cat No. KTSDEHP) and cultured according to the company’s protocol on grids at 30,000 to 40,000 cells/cm^2^ to either DIV 1 or 2 before being picked up with forceps and loaded on the Vitrobot. Grids were hand-blotted through the side port, from the back, using forceps holding blotting paper for 2 seconds. After plunge-freezing, samples were stored in liquid nitrogen until they were loaded into the cryo-TEM for tilt series collection.

### Specimen preparation, Image Collection and Reconstruction

Stock actin (10 mg/mL) was diluted 1:10 into ATP-supplemented general actin buffer from Cytoskeleton Inc. (Cat # BSA01-001, 5mM Tris-HCl pH 8.0, 0.2 mM CaCl^2^, and 0.2 mM ATP), and left on ice for 30 min. Next, 10 uL of actin polymerization buffer (Cat # BSA02-001, 50 mM KCl, 20 mM MgCl^2^, and 1 mM ATP) was added, mixed well, and placed at RT for one hour to polymerize. After polymerization, 2.5 uL of Cofilin (5 mg/mL) was added to a tube and diluted in 10 uL of 20 mM Tris-HCl 6.6 Buffer, before adding 12.5 uL of polymerized actin and letting it sit for 30 min at RT. Before vitrifying CaMKII, stock aliquots dialyzed in 10 mM HEPES, pH 7.4, 0.1 mM EGTA, 200 mM KCl, and 20% glycerol, were thawed and diluted from 3.4 mg/mL to 1 mg/mL in dialysis buffer. Isolated chromatin samples were incubated for 20 minutes with MgCl2 and then mixed with a suspension of 10 nm fiduciary gold particles, as described fully in ^28^.

All samples were plunge frozen on R2/2 200 mesh Quantifoil EM grids, using a Vitrobot Mark IV (Thermo Fisher). Copper grids were used for purified particles in suspension and gold grids were used for culturing neurons. For purified actin/cofilactin, chromatin and alpha-CaMKII, 3 µL of sample were applied to freshly glow-discharged grids and blotted for 3.5 s with force 5.

Tilt series were collected on a Titan Krios G3 or G3i (Thermo Fisher) 300 kV cryo-TEM, equipped with a Bioquantum energy filter (Gatan) and either a K2 or K3 direct electron detector, respectively. Using Tomography 5 (Thermo Fisher), data was collected from -60 to 60 degrees in either a dose-symmetric scheme or a split scheme with 2 degree increments for everything except purified chromatin, which was imaged with 5 degree tilt increments. Total electron dose was maintained between 60-180 e^-^/Å^2^, and defocus was maintained at ∼-5 microns except as otherwise stated. A range of magnifications were used, depending on the sample, from 6.6 - 20 Å/pix after binning. Tomographic alignment and reconstruction was carried out using IMOD. All tomograms were reconstructed with weighted back projection and were binned 4x before segmentation.

### PDB structures

See Table S1 for PDBs used and a description of their modifications.

### Generating Synthetic Training Data

#### Modeling Parameters

To build synthetic cytoplasmic models we used the following parameters:

Pixel Size: 6.6 - 14 Å/pix

Volume Dimensions: 400x400x50 (XYZ)

Particle Density: 0.8

Layers: 4 (targets, large and small; distractors, large and small)

Iterations: 1200 (large targets), 1200 (small targets), 2400 (large distractors), 4800 (small distractors)

Membranes: 10

Constraint: Sides

#### Simulation Parameters

To simulate tilt series for reconstruction we used the following parameters:

Tilt Parameters: +60 to -60 with two, three, or five degree tilts

CTF Parameters: Defocus = -6uM, Voltage = 300 kV, Aberration = 2.7 mm, Envelope = 1

Imaging Parameters: Total Dose = 70-150 e^-^/Å^2^, Radiation Damage = 1, Tilt error = 0

### Deep Learning and Performance Testing

All U-Net building, training, inference, and performance testing (DICE scores calculations) was done within the Dragonfly software suite by Comet, Inc. Non-commercial licenses are available through their website at (https://dragonfly.comet.tech/), and an open-source tutorial video can be found at www.jove.com/t/64435/deep-learning-based-segmentation-of-cryo-electron-tomograms.

In all cases, 11-slice 2.5D U-Nets were used with the following hyperparameters: Patch Size = 64 or 128 pixels^2^, Stride Ratio = 0.5 or 1, Batch Size = 8, Epochs = 100-300. The loss function used for semantic segmentation was “CategoricalCrossEntropy” and the optimization algorithm was “adadelta”.

### Quantifying Growth Cone Segmentations

After segmentation, quantifications of individual proteins of interest involved a few different methods. Microtubules were individually counted by eye. Ribosomes were estimated by splitting unconnected components within the class into individual labels. The TriC class was estimated the same as ribosomes but was further separated with a watershed transform to generate centroid coordinates. Actin and cofilactin filaments were quantified by total voxel count within each class.

### TriC cluster characterization

Clustering analysis of TriC particles was performed with the DBSCAN algorithm ^38^, using the implementation in MATLAB. Briefly, clusters are iteratively calculated from points that have a minimum number of neighbors, and are then merged in subsequent iterations if they are in close proximity. The key parameter, epsilon, was set at 50 nm for the clustering process, meaning any particles within 50 nm of the seed point would be considered clustered. Nearest Neighbor analysis within clusters of TriC was performed within MATLAB using a custom script.

### Training NeuralSeg

Synthetic tomograms (400×400×50) with varying pixel sizes (12-14Å/pixel), doses (70-150 e^-^/Å^2^), and defocus values (-4 to -6 µM) were generated using a custom CTS Matlab script, along with their corresponding ground truths, and concatenated into two blocks of synthetic tomograms and atlases via a Python script for all iterations of training NeuralSeg (Figure S7A). The blocks contained 7 classes: membrane, 5 proteins of interest (microtubules, actin, cofilactin, ribosomes, and TriC) and background containing large and small distractor molecules from a randomized PDB pool (table S1). Initial real data included a cropped neuronal tomogram (400×400×14) and its hand-segmented ground truth that had the seven classes listed (Figure S7B 1.1).

A base network was co-trained on the initial real data and the synthetic blocks and inferred to a set of ten neuronal tomograms. Regions where the network failed to segment correctly, e.g., cofilactin bundles, overlapping microtubules, and mitochondrial membranes, were identified (Figure S7B 1.2), cropped to 400×400×Z where Z = 10–50, hand-corrected, and concatenated via Python with the initial real data to make two updated blocks of real tomograms and atlases. Second iteration of NeuralSeg was trained on described blocks of synthetic and real data and then inferred to three non-neuronal tomograms (EMDB-19370, EMDB-15989, and EMDB-27479). Regions where the network misclassified, e.g., multi-layered membranes in eukaryotic cells and bacterial double membranes, were processed the same as in the first iteration to update data for retraining (Figure S7B 2.1-2.3). The final iterations included regions consisting of common tomographic artifacts (e.g., ice-surface contamination, fiducial gold, carbon edges, and fringing) annotated as background (Figure S7B 3.1-3.4). All data blocks were z-score normalized before concatenation and import to Dragonfly.

### Software Download

CTS can be downloaded from its [Github](https://github.com/carsonpurnell/cryotomosim_CTS) page. The github project also contains installation instructions, required software, and preliminary tutorials on its use and application.

## Acknowledgements

This study was supported by the Penn State College of Medicine and the Department of Biochemistry and Molecular Biology, as well as Tobacco Settlement Fund (TSF) grant 4100079742 (to M.T.S), the National Institutes of Health (NIH) grant NS126448-01A1 (to M.T.S), NSF grant MCB-1817929 (to F.A.H.), NSF Grant CHE-220412 (to F.A.H and M.N.W.), NIH Grant R01GM138887 (to F.A.H and M.N.W.), and NSF grant 1911940 (to S.G.). M.N.W. acknowledges the William Wheless III Professorship.

The CryoEM and CryoET Core (RRID:SCR_021178) services and instruments used in this project were funded, in part, by the Pennsylvania State University College of Medicine *via* the Office of the Vice Dean of Research and Graduate Students and the Pennsylvania Department of Health using Tobacco Settlement Funds (CURE). The content is solely the responsibility of the authors and does not necessarily represent the official views of the University or College of Medicine. The Pennsylvania Department of Health specifically disclaims responsibility for any analyses, interpretations, or conclusions.

---

**Figure S1:**
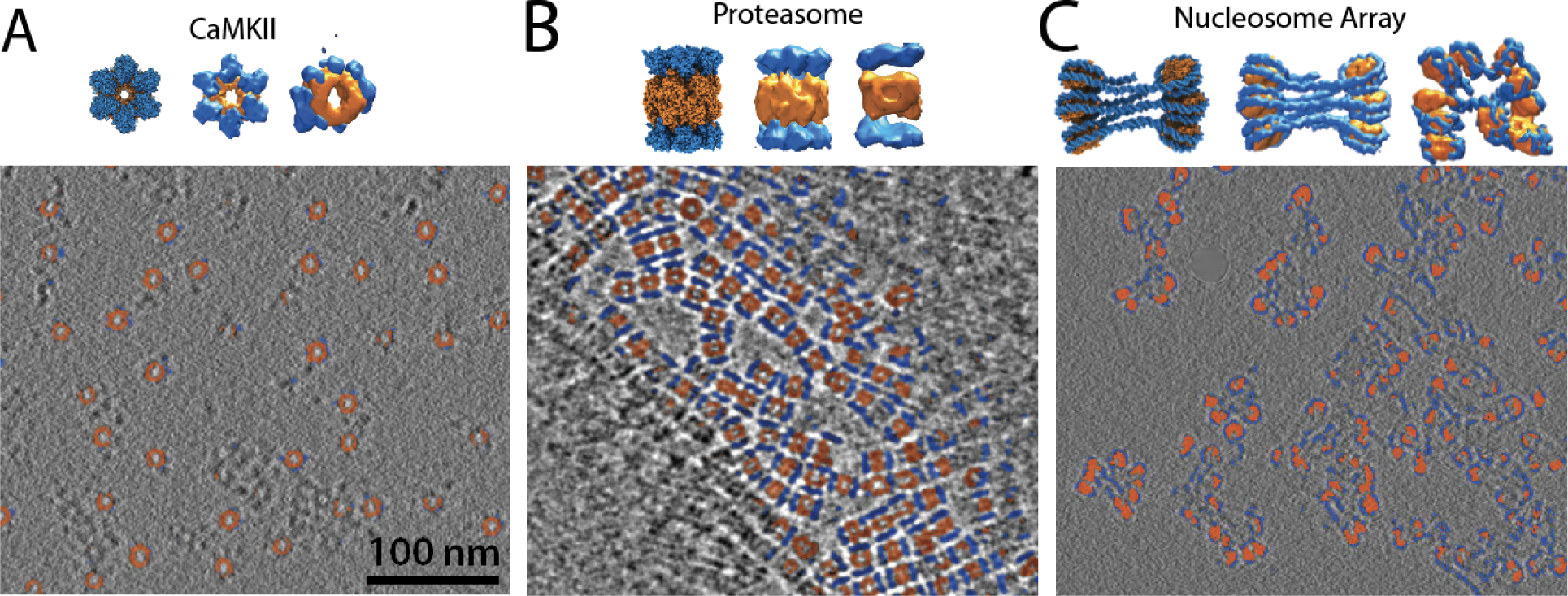
Sub-molecular segmentation using synthetic data. (A) The top panel contains a space-filling atomic model of CaMKII (PDB 3SOA), that model rendered as a colorized density map, and a 3D rendering of a segmented molecule from the corresponding tomogram in the lower panel (EMD-41263) showing segmentation of the core (orange) and catalytic domains (blue). (B) Similar panels showing the approaches effectiveness in segmenting the outer and inner rings of purified proteasomes in EMD-7152 (PDB 5FMG). (C) Similar panels showing the approaches effectiveness in segmenting DNA and histones within purified nucleosome arrays (PDB 6HKT).

**Figure S2:**
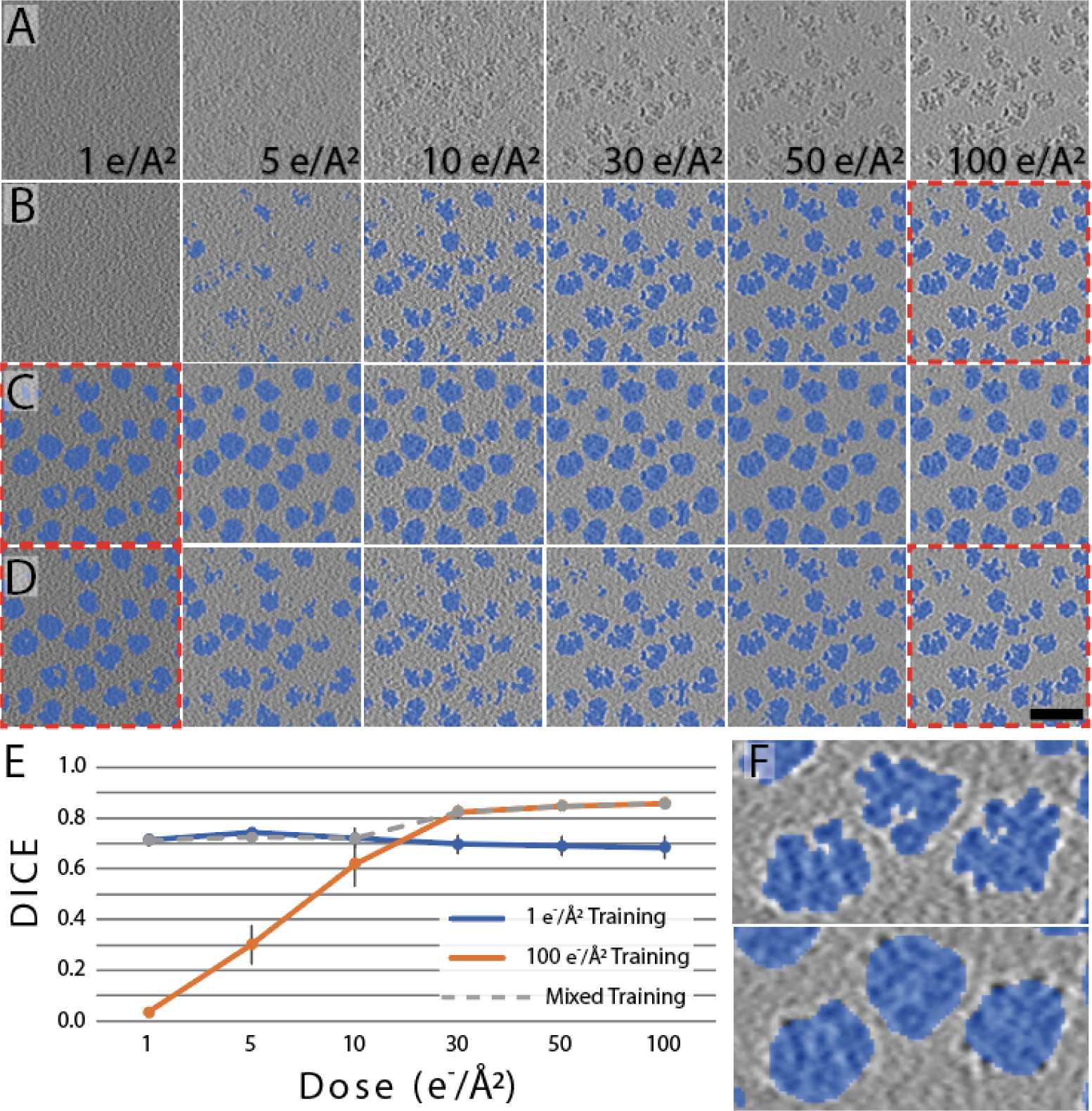
The effect of electron dose on training segmentation networks. (A) A series of synthetic tomograms generated from the same model (randomly oriented ribosomes), but with different electron dose (-1). (B) Ribosomal segmentations (blue) generated from a single network trained on the highest-dose dataset (red-dashed box). (C) Ribosomal segmentations generated from a single network trained on the lowest-dose dataset (red-dashed box). (D) Ribosomal segmentations generated from a single network trained on both the highest- and lowest-dose datasets (red-dashed boxes). (E) Line graph of the DICE scores for each segmentation compared in terms of electron dose. (F) Close-up views of segmentations from 100 e^-^/Å^2^ simulations in (C) and (D), Top and Bottom, respectively.

**Figure S3:**
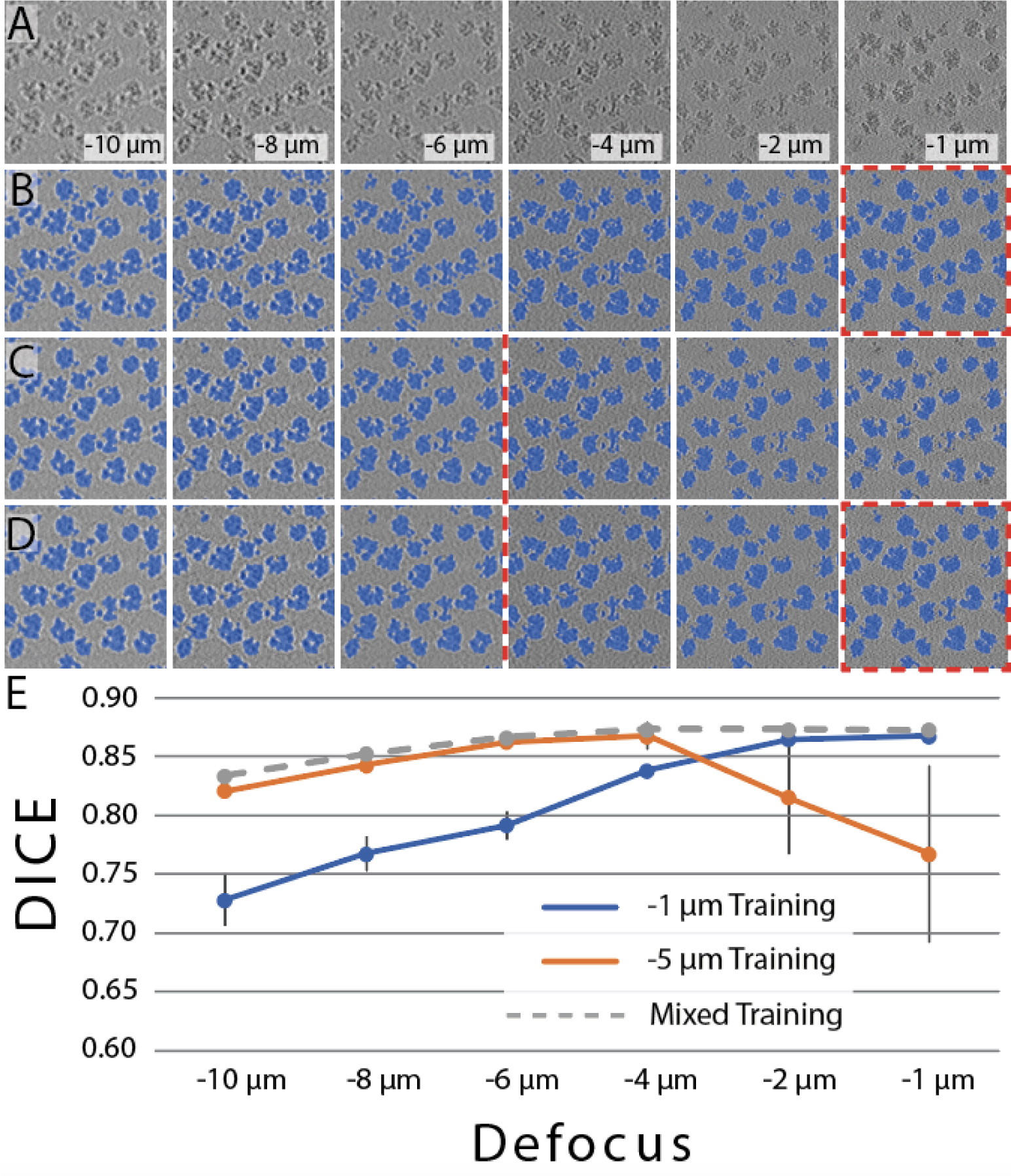
The effect of defocus on training segmentation networks. (A) A series of synthetic tomograms generated from the same model (randomly oriented ribosomes), but with different defocus values. (B) Ribosomal segmentations (blue) generated from a single network trained on the -1 µM defocus dataset (red-dashed box). (C) Ribosomal segmentations generated from a single network trained on a -5 µM defocus dataset (red-dashed line). (D) Ribosomal segmentations generated from a single network trained on both the -1 and -5 µM defocus datasets (red-dashed line and box). (E) Line graph of the DICE scores for each segmentation compared in terms of image defocus.

**Figure S4:**
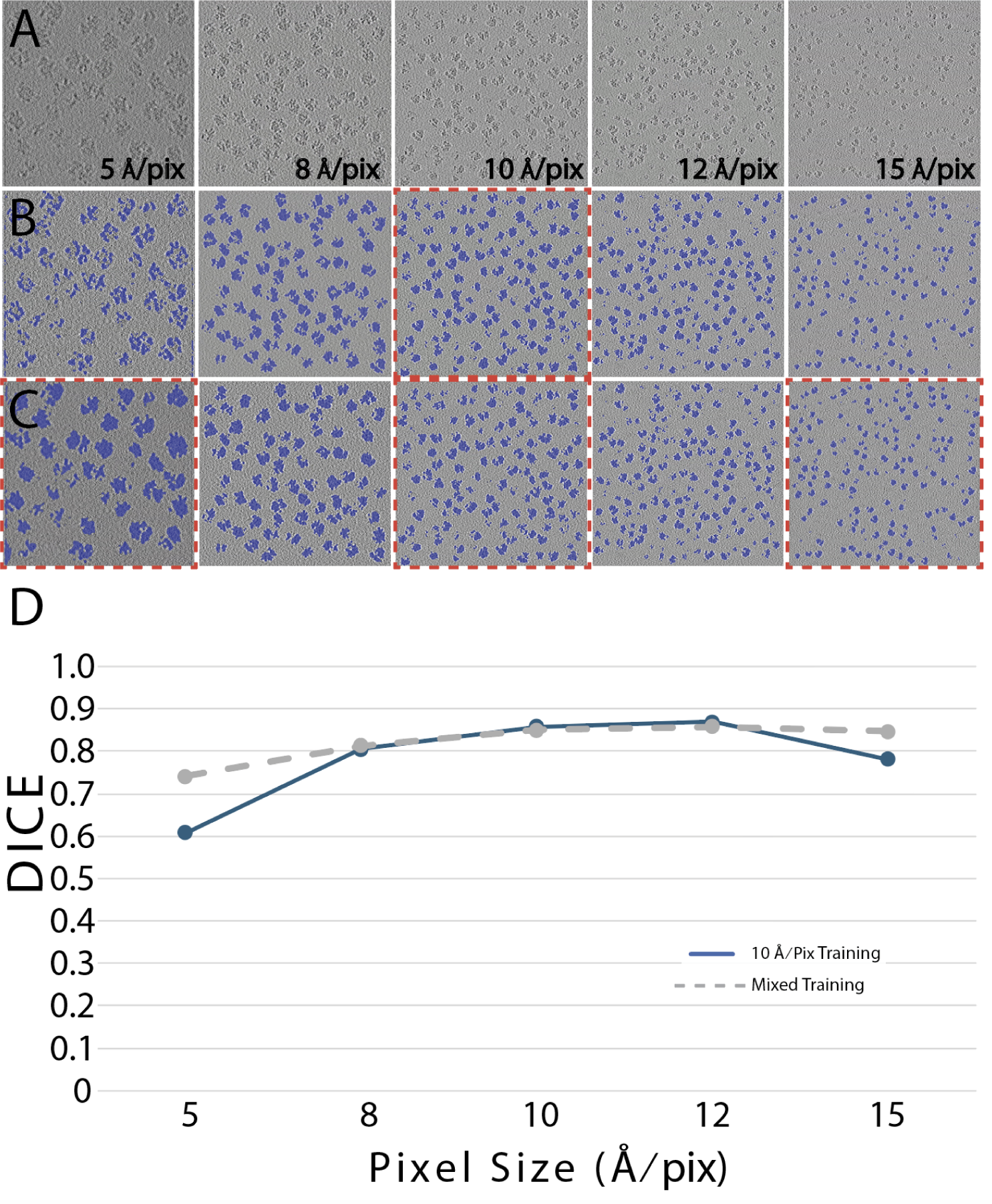
The effect of Pixel Size on training segmentation networks. (A) A series of synthetic tomograms generated from models of randomly oriented ribosomes along a gradient of pixel sizes (5-15 ang/pix). (B) Ribosomal segmentations (blue) generated from a single network trained at 10 ang/pix (red-dashed box). (C) Ribosomal segmentations generated from a single network trained at 5, 10, and 15 ang/pix (red-dashed boxes). (D) Line graph of the DICE scores for each segmentation compared in terms of pixel size.

**Figure S5:**
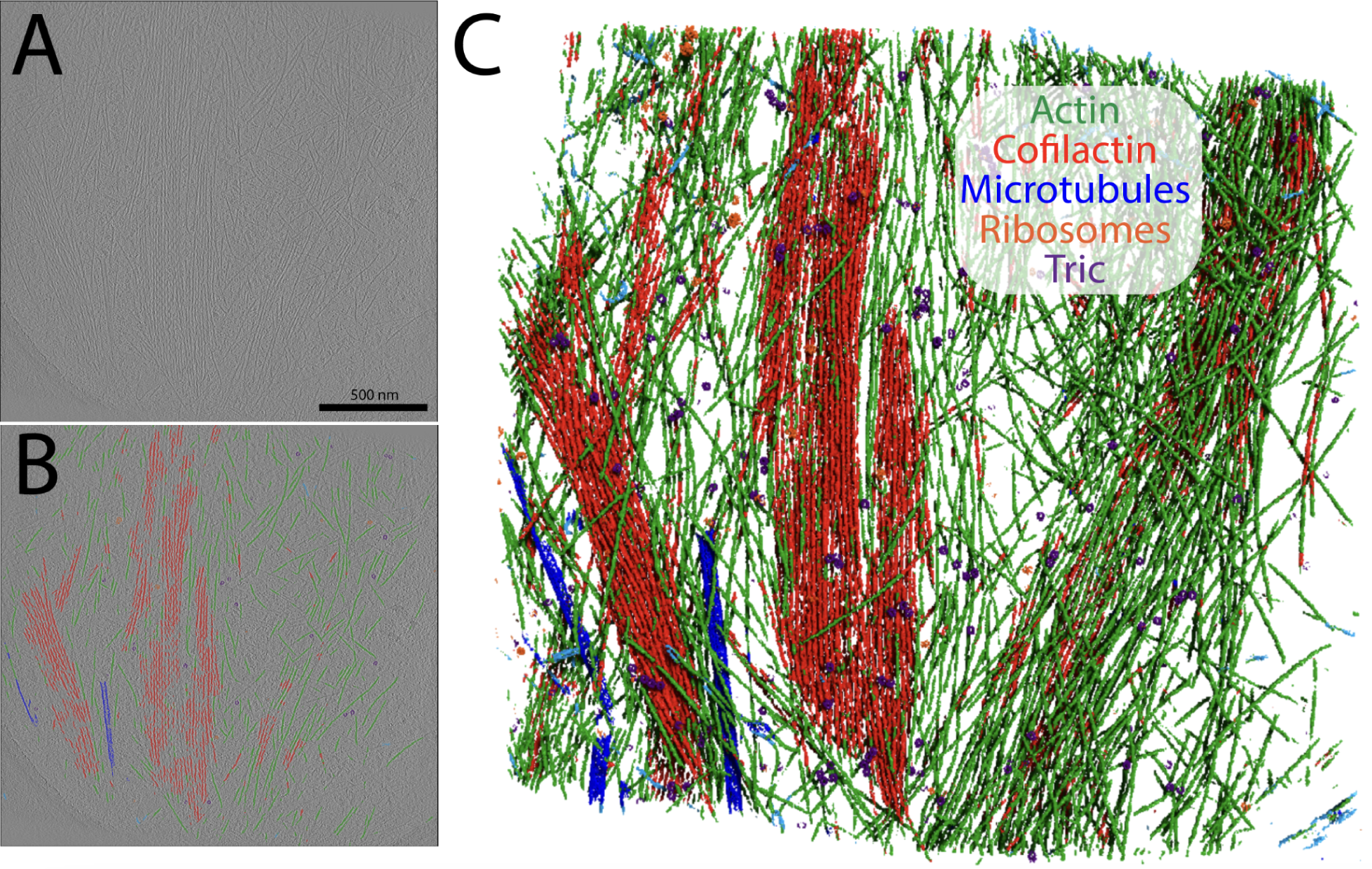
Co-trained networks easily differentiate between actin and cofilactin in situ. (A) Slice from a cryotomogram taken within the transition zone of a neuronal growth cone, and (B) its segmentation by a co-trained UNet. (C) High-resolution 3D segmentation of the tomogram in (A), where bare actin and cofilactin are distinguished either individually, as bundles, or mixed bundles.

**Figure S6:**
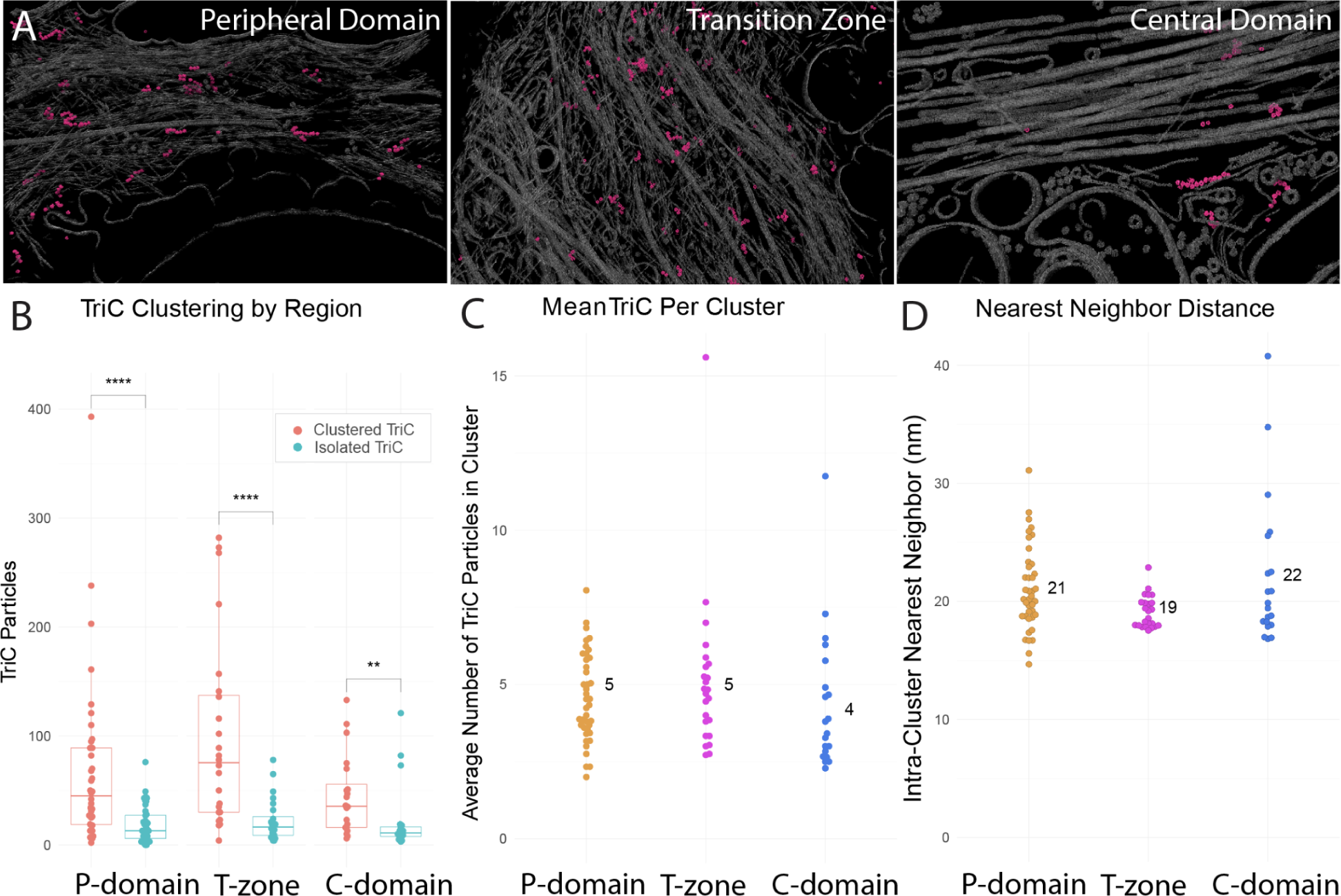
TriC clustering in growth cones. (A) Representative segmentations of TriC clusters (pink) in tomograms from each domain of the growth cone. (B) Distribution of isolated and clustered TriC across domains. (C) Mean number of TriC particles in each cluster across domains. (D) Measured intra-cluster nearest neighbor distance for TriC.

**Figure S7:**
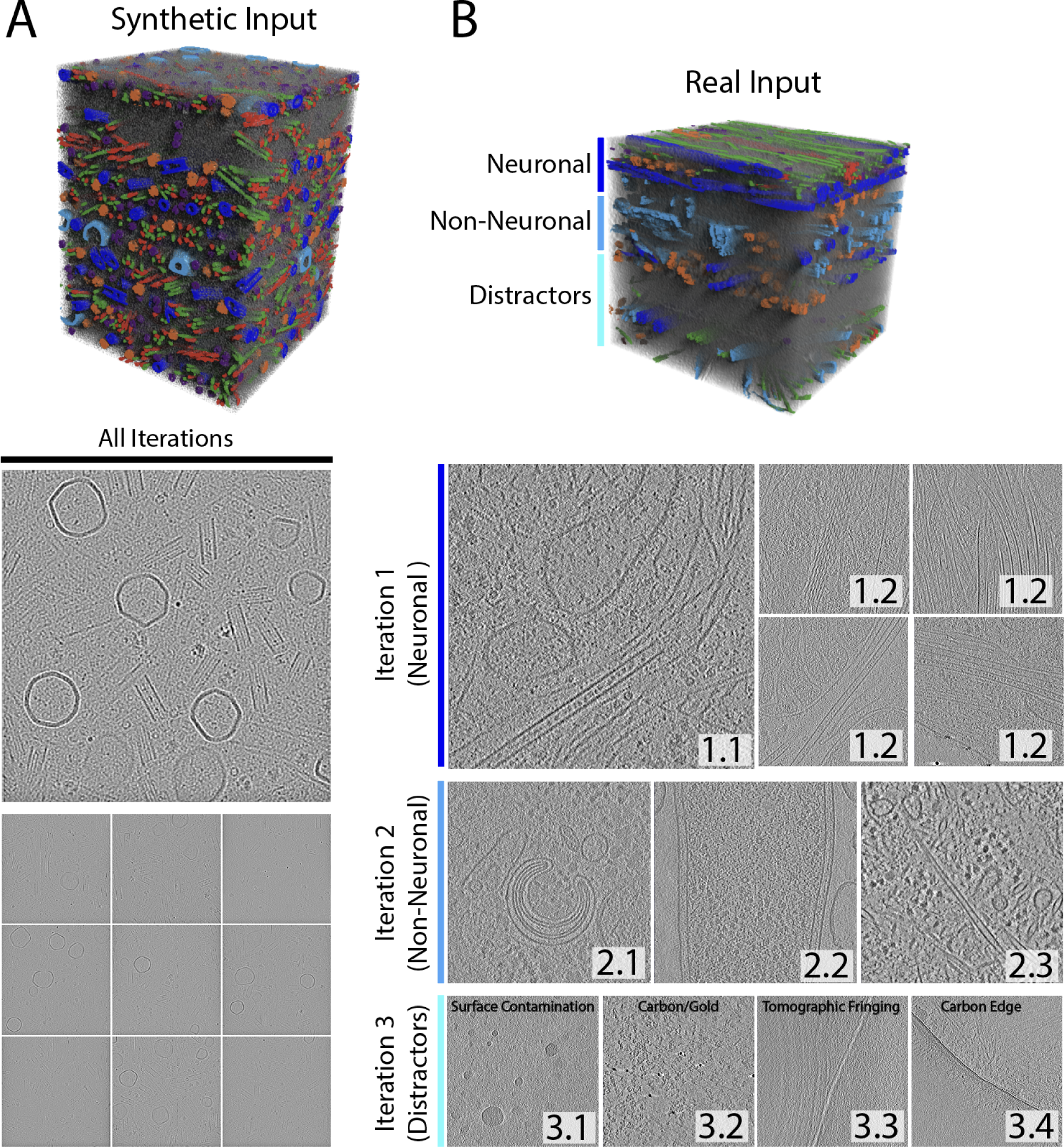
Training inputs for the development of NeuroSeg. (A) Block of concatenated synthetic tomographic data. All synthetic tomograms are 400x400x50 voxels and generated programmatically by CTS to vary randomly within a set range of both modeling and simulation parameters. In the lower panel, individual synthetic tomograms are displayed. They contain the primary targets (membrane, microtubules, actin, cofilactin, ribosomes and TriC) as well as randomized sets of molecules from a pool of distractor PDBs. (B) Block of concatenated real tomographic data. All real tomograms are 400x400xZ voxels, where Z varies between 10 and 50. During each iteration of the network, the synthetic data was used as a base for co-training with the real inputs. For each iteration, segmentations based on the previous round of training were used as a starting point for cropping and class remapping, as well as repair of false positives/negatives. Hand-corrected patches of data were then fed back into the next iteration of training.

**Figure S8:**
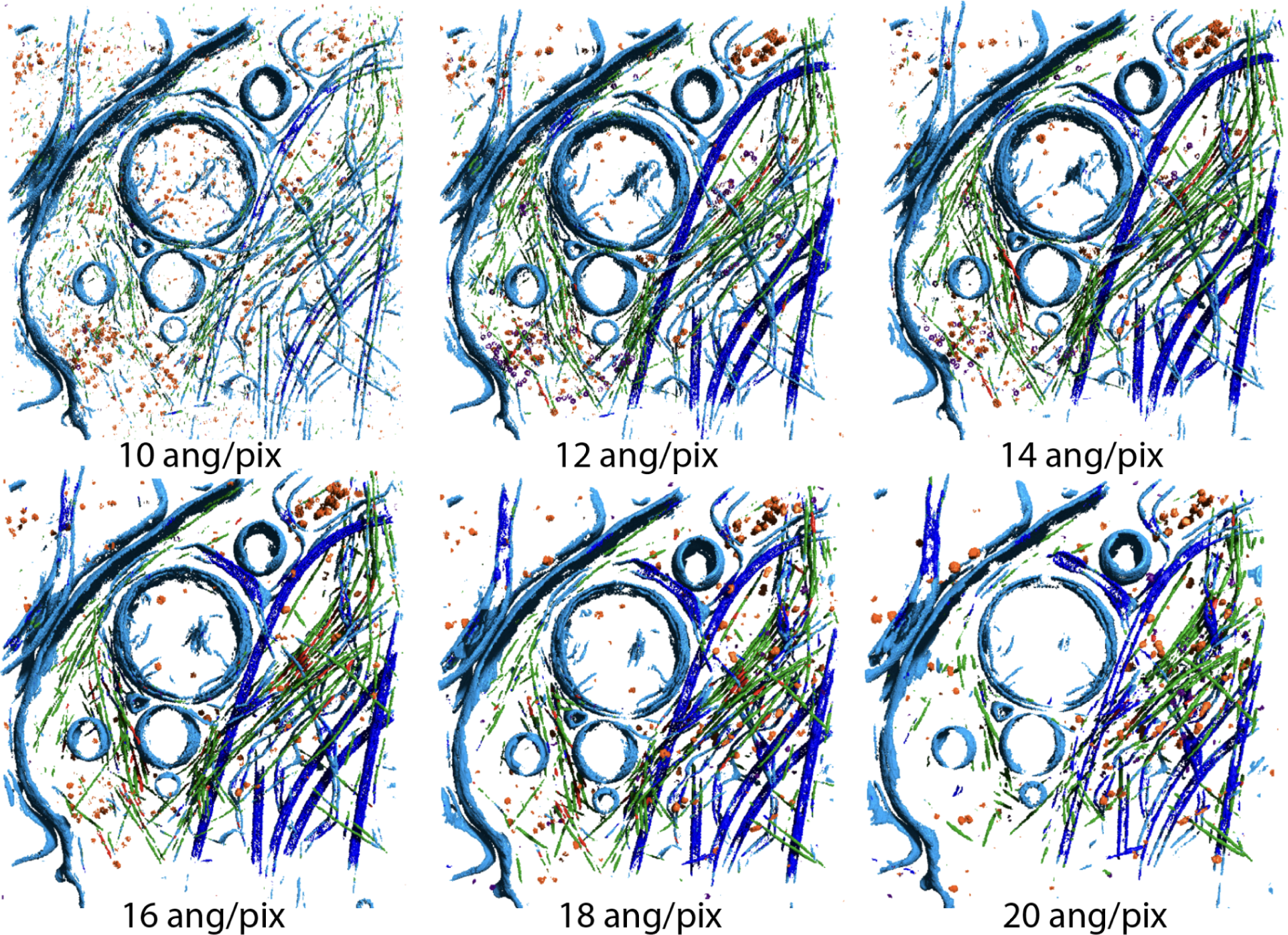
NeuralSeg works across a range of pixel sizes. At the top left is the segmentation of the original tomogram reconstructed at 10 angstrom/pixel, followed by a series of segmentations of the same tomogram after rescaling in 20% increments to 20 ang/pix.

**Figure S9:**
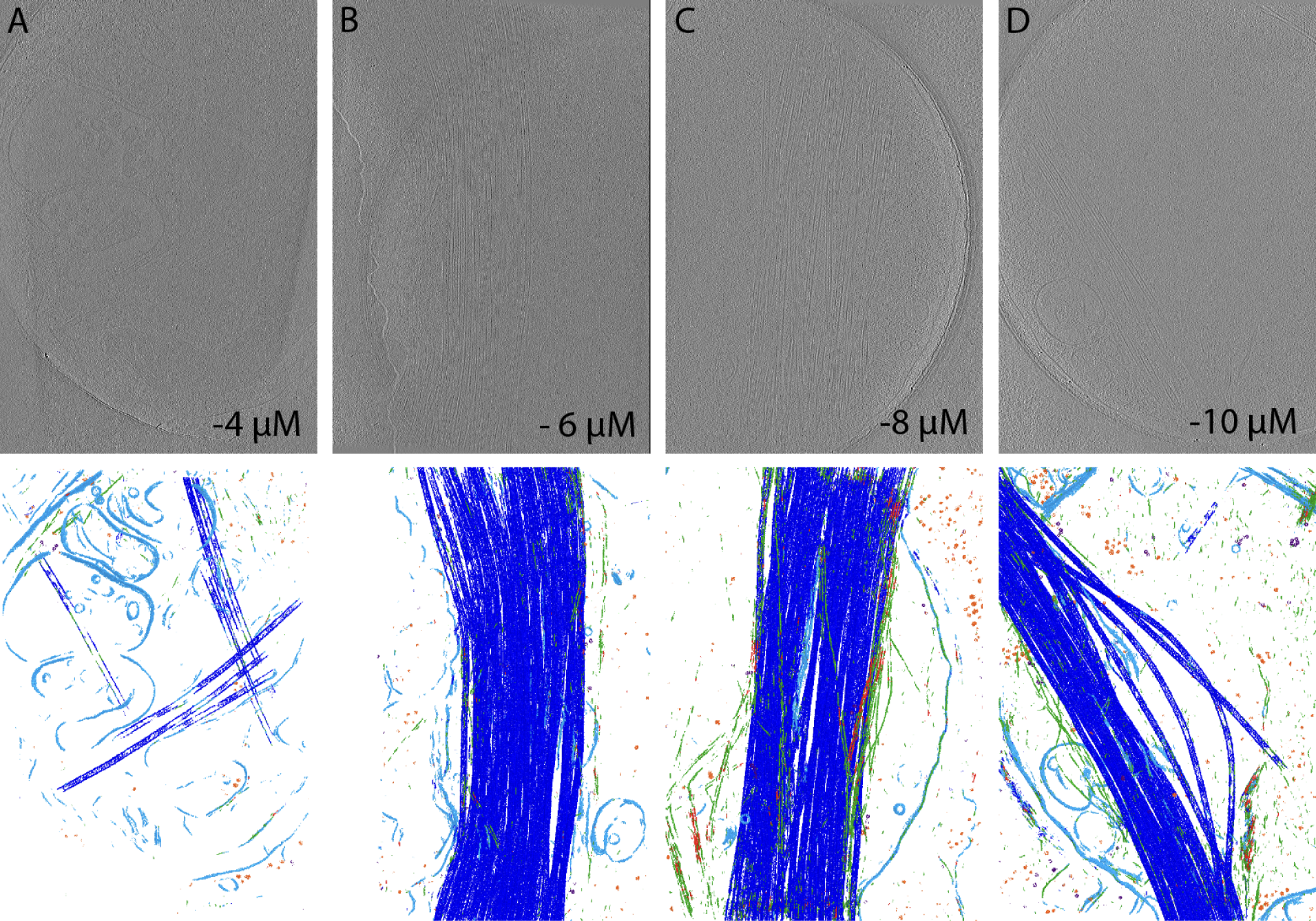
NeuralSeg works across the range of defocus from -4 to -10 µM. Top panels are slices through neuronal tomograms collected at different defoci, from -4 to -10 µM, and below are their corresponding segmentations by NeuralSeg.

**Figure S10:**
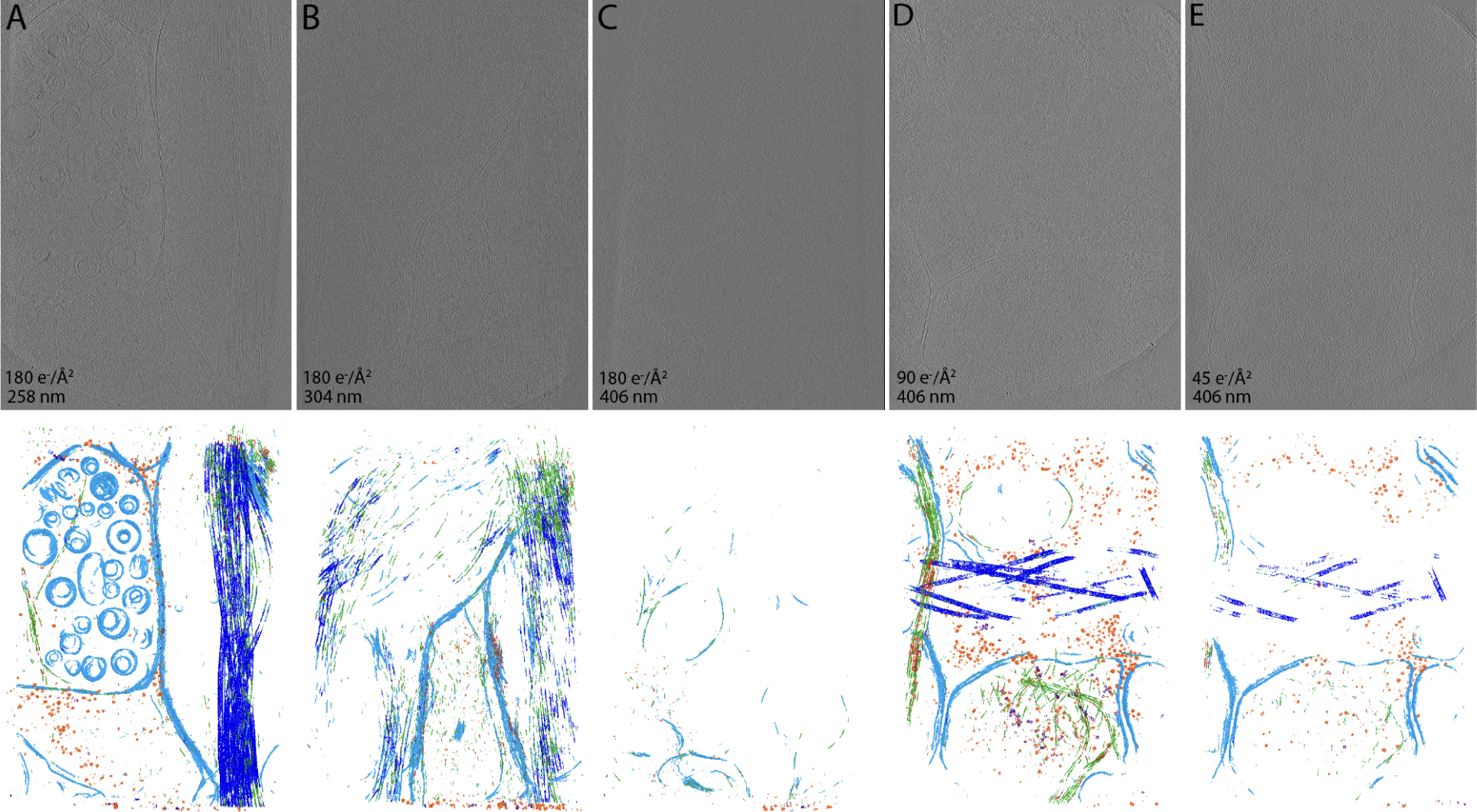
Tomogram contrast resulting from thickness and electron dose affects NeuralSeg’s performance. Top panels are slices through six tomograms and their corresponding segmentations by NeuralSeg on the bottom. (A) through (C) are tomograms collected at the same electron dose of 180 e^-^/Å^2^ but have increasing thickness from 258 nm to 406 nm. (D) and (E) are tomograms of the same position, and therefore the same thickness of 155 nm, but at different electron doses, 90 e^-^/Å^2^ and 45 e^-^/Å^2^.

**Table S1:** PDBs and their modifications for simulation.

| Protein | PDB(s) | Modifications |
| --- | --- | --- |
| Actin | 6t1y | Elongated helically to 39 subunits. |
| Cofilactin | 3j0s | Elongated helically to 33 subunits. |
| Microtubules | 6o2t | Elongated helically by 2x, filled with 3 MIPS. |
| CaMKII | 3soa,<br>5u6y | Split into catalytic and association domains. |
| Proteasome | 5fmg | Split into alpha and beta component submodels. |
| Chromatin Array | 6hkt | Spit into DNA and histones |
| Ribosome | 6ks8 |  |
| TriC | 4ujd |  |
| Distractors |  |  |
| Small (<200 kDa) | 2q0u, 1sjj, 1qtx, 2cg9, 4uic, 5csa, 6vgr, 7e6g, 7wbt, 1exr |  |
| Large (>200 kDa) | 1qvr, 2dfs, 3soa, 5fmg, 5o32, 6igc, 6ksp, 6u8q, 7bkc, 7nhs |  |

## CTS Overview

### Modeling Approach

CTS models are essentially Coulomb potential maps based on atomic structure files (.pdb and .cif) with limited support for models based on density maps. CTS follows the weak phase object approximation, which is sufficient for most cryoEM specimens. Models are generated primarily using brute force, with individual structures tested at potential insertion sites several times, repeated over many iterations. Use of efficient code allows CTS to generate models quickly, with small models (∼400x400x100 px) taking only a few minutes to create depending on density. CTS has separate functions for generating full atomic models and coarse voxelwise models depending on computational resources, resolution, and use case.

### Modeling steps

A CTS model starts with any of the standard features opted into: a carbon support film, vesicle membranes, and boundary constraints. Input structures are then added in a series of layers set by the user, allowing precise control over relative abundance and occupancy of different inputs. Next, gold fiducial beads can be added, and the model is embedded in vitreous ice. Finally, an atlas is generated with each voxel labeled with the class of its main component alongside a simulation run, matched to its final size.

### Modeling features

Clustering: structures can be placed as local clusters isotropically, or in packed bundles to model filaments like actin.

Complexes: structures can be subdivided, with each subdivision labeled separately in the output atlas. This allows subdomains of very large proteins or different proteins in a complex to be distinguished. A variant ‘assembly’ allows randomized occupancy of subdivisions for easier implementation of nonuniform structures.

Membrane embedding: structures can be placed in the membrane as a transmembrane or membrane-associated protein. The OPM database provides compatible structure files, but CTS has a tutorial for adapting any structure for use in this way with UCSF Chimera.

Vesicle placement: structures can be placed exclusively inside membrane-bound vesicles or exclusively outside, not merely globally random locations.

### Simulation approach

CTS also uses a coarse-grained method to generate simulations from input models, operating on bulk interactions at a pixel level rather than atomic and wave interactions. This enables fast runtimes on minimal hardware, without sacrificing utility for our deep learning purposes. CTS has a large number of input parameters whose default values cover modern hardware but can cover a range of possibilities from standard to reasonable and even technically impossible imaging capabilities. Primary control parameters are the same as standard imaging parameters: microscope characteristics and imaging factors including tilt increment and limits, defocus, electron dose, and tilt scheme. Advanced options include control over radiation damage, deviation from target tilt angles, inelastic electron scattering, and generation of “ideal” images that lack CTF modulation and probabilistic dose sampling. CTS has two simulator functions: the first a purely volume-based implementation that uses a 2.5d approximation of defocus and CTF, and a more intensive option that requires an atomic model but implements a fully 3d CTF at the cost of runtime.

### Simulation Components

#### Tilt Projection

For the volumetric method, CTS currently relies on the external IMOD^24^ xyzproj command to create the initial model projections. Regardless of method, CTS can project tilts around either the X or Y axis and though it expects a balanced tilt series can project any arbitrary set of angles in any order. This allows single micrographs, multiframe tilts, and complex acquisition series. CTS also provides a parameter for tilt error, which introduces a scaling random variance to the true angle that is projected from the target angle.

#### Dose sampling

The density contribution to contrast is simulated by sampling electrons from a distribution of the tilt angle’s scattering potential (Fig. 1D). For each tilt, the camera-detected dose is adjusted based on the maximal DQE of the detector as well as inelastic scattering of electrons away from the path of the detector. This dose-adjusted scattering map is used as the lambda parameter of a poisson distribution from which detected electrons are drawn. The following equation is used to determine transmitted dose:

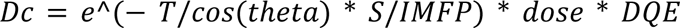

Dc is the corrected dose transmitted, S is the scattering factor (1), IMFP is the inelastic mean free path of vitreous ice (3.8nm), T is the thickness of the sample, theta is the tilt angle, and DQE is the detector quantum efficiency of the camera.

#### CTF intensity modulation

Once tilt images are projected, they are modulated by a contrast transfer function (CTF) based on the pixel size and defocus, as well as microscope parameters such as the accelerating voltage and spherical aberration (Figure 1E). As mentioned previously only with the atomic simulator is CTF modulation fully 3d, while the volumetric simulator uses a 2.5D method with the defocus approximated in a series of overlapping 2D strips parallel to the tilt axis.

The CTF in fourier space is computed according to the following functions:

1. *CTF* = *E* * ((1 − *Q*)*sin*(*eq*) + *Qcos*(*eq*))
2. *eq* = *pi*/2 * (*CS* * *L*^3^ * *k*^4^ − 2 * *Dz* * *L* * *k*^2^)
3. *E* =*e*^−(*k*/(*sigma*\**nyquist*))^2^^
4. *L* = *H* * *c*/*sqrt*(*e* * *V* * (2 * *m* * *c*^2^ + *e* * *V*))

Overall equation for the CTF profile (1), wave component equation (2), envelope function of the contrast (3), and the calculation of the relativistic electron wavelength (4).

Where L is the relativistic electron wavelength, CS is spherical aberration, K is the spatial frequency, Dz is defocus, sigma is the envelope factor (.9), and Q is the amplitude contrast factor (.07). In 4, H is the planck constant, e the electron charge, c the speed of light, m the electron mass, and V the acceleration voltage.

#### Radiation damage

Radiation damage is modeled in a very simplified fashion. Before electrons are sampled for each tilt angle, that tilt projection is corrupted by two operations scaled by the cumulative electron dose transmitted. The first operation is a smoothing step that reduces signal clarity in higher-resolution images, and the second is a layer of gaussian noise applied to the whole tilt image.

#### Tomographic reconstruction

CTS outputs are ready for immediate reconstruction without any other processing necessary. CTS performs reconstruction of the simulated tilt series (Fig. 1F) using IMOD’s tilt command with the target tilt angles, followed by using the IMOD command trimvol to rotate the tomographic reconstruction to a standard orientation. The lack of CTF correction (and alignment, in the case of introduced alignment error) are deliberate, as the baseline values for each contribute more to very small scale contrast and signal, rather than large-scale errors. These correspond to higher quality data that similarly would not need further refinement.

